# From iron to antibiotics: Identification of conserved bacterial-fungal interactions across diverse partners

**DOI:** 10.1101/2020.03.19.999193

**Authors:** Emily C. Pierce, Manon Morin, Jessica C. Little, Roland B. Liu, Joanna Tannous, Nancy P. Keller, Benjamin E. Wolfe, Kit Pogliano, Laura M. Sanchez, Rachel J. Dutton

## Abstract

Microbial interactions are major determinants in shaping microbiome structure and function. Although fungi are found across diverse microbiomes, the mechanisms through which fungi interact with other species remain largely uncharacterized. In this work, we explore the diversity of ways in which fungi can impact bacteria by characterizing interaction mechanisms across 16 different bacterial-fungal pairs, involving 8 different fungi and 2 bacteria (*Escherichia coli* and *Pseudomonas psychrophila*). Using random barcode transposon-site sequencing (RB-TnSeq), we identified a large number of bacterial genes and pathways important in fungal interaction contexts. Within each interaction, fungal partners elicit both antagonistic and beneficial effects. Using a panel of phylogenetically diverse fungi allowed us to identify interactions that were conserved across all species. Our data show that all fungi modulate the availability of iron and biotin, suggesting that these may represent conserved bacterial-fungal interactions. Several fungi also appear to produce previously uncharacterized antibiotic compounds. Generating a mutant in a master regulator of fungal secondary metabolite production showed that fungal metabolites are key shapers of bacterial fitness profiles during interactions. This work demonstrates a diversity of mechanisms through which fungi are able to interact with bacterial species. In addition to many species-specific effects, there appear to be conserved interaction mechanisms which may be important across microbiomes.

## INTRODUCTION

Despite awareness that fungi have an immense capacity to produce biologically active metabolites and to reshape ecosystems, fungi are frequently overlooked in microbiome studies in favor of a bacterial-centric focus^1–3^. Recently, fungi and other microeukaryotes have received increased attention in sequencing-based studies^4–9^, and there is growing interest in exploring the role fungi and bacterial-fungal interactions play in environmental and host-associated microbiomes^10–17^. However, the broader patterns and diversity of bacterial-fungal interaction mechanisms have been challenging to characterize given the abiotic and biotic complexity of many microbiomes. While specific interaction mechanisms have been elucidated for particular pairwise bacterial-fungal associations, little is known about the conservation of these mechanisms for other bacterial-fungal pairs and thus about general and widespread interaction mechanisms that could be a major factor in shaping microbiomes.

Fermented foods have been developed as experimentally tractable systems to study multikingdom microbial communities^18–20^. Cheese rind biofilms, in particular, represent an ideal system to investigate the diversity and conservation of bacterial-fungal interactions due to the presence of culturable bacterial and fungal species representing diverse phyla. Previous work has shown that biofilms that form on the surface of cheese are composed of an average of six bacterial and three fungal genera^18^. These genera are frequently found in natural and human environments, suggesting that interactions in this system may resemble interspecies interactions in other microbiomes. Communities of isolated fungal and bacterial species from cheese can be reassembled to investigate how these members interact in co-culture^21^. Prior work in this system has demonstrated that fungi in rind biofilms can have both strong positive and negative impacts on bacterial growth and that specialized metabolites can impact these interactions^18, 19, 22, 23^.

To systematically characterize the potential effects of fungal species on bacteria within this system, we have employed an interdisciplinary approach. Specifically, we used the high-throughput genetic screen RB-TnSeq^24^, RNA-Seq, bacterial cytological profiling, and metabolomics to investigate the diversity of mechanisms through which eight diverse fungal species impact two bacteria *(Escherichia coli* or a cheese-associated *Pseudomonas psychrophila).* A previous study used RB-TnSeq in the cheese rind microbiome system to identify interspecies interactions and to characterize the occurrence of higher-order interactions in a microbial community^21^. RB-TnSeq has been used in other systems to identify genes important for host colonization, to determine essential gene sets, and to characterize genes of unknown function^25–27^.

Here, to infer potential mechanisms of bacterial-fungal interactions, we have developed customized experimental and computational RB-TnSeq pipelines to identify genes impacting bacterial fitness in a mixed biofilm with a fungal partner. We observed a diversity of gene functions that affect bacterial fitness in the presence of a fungal partner. Additionally, we found conservation among the impacts of yeasts and filamentous molds on bacteria; a key example of this is the widespread effect on iron metabolism in bacteria, which is mediated by the acquisition of fungal siderophores. Furthermore, we observed similar effects when we expanded our analysis to include soil and skin fungi, suggesting that this mechanism is relevant not only within cheese rind biofilms, but also in other systems. Consistent with the understanding that fungal species have diverse metabolic repertoires, we find several examples of species-specific effects including the apparent production of antimicrobial compounds by filamentous fungal species. Analysis of a fungal mutant defective in secondary metabolite production revealed changes in the bacterial response compared to the wild type (WT) strain, including genes involved in siderophore uptake and antibiotic resistance. This work provides new perspectives on the biology and mechanisms of bacterial-fungal interactions and highlights the key roles of fungi in microbiomes.

## RESULTS

### Selection of Bacterial-Fungal Interaction Partners

To represent fungal diversity within the cheese environment, we selected a panel of 8 species of commonly found yeasts and filamentous fungi (Figure 1). The selected fungal species from cheese rind microbiomes, which represent five different genera and three fungal classes, include two yeasts, *Candida* sp. str. 135E and *Debaryomyces* sp. str. 135B, and filamentous fungi *Penicillium* sp. str. #12, *Penicillium* sp. str. SAM3, *Penicillium* sp. str. RS17, *Fusarium* sp. str. 554A, *Scopulariopsis* sp. str. JB370, and *Scopulariopsis* sp. str. 165-5. These fungal genera were all present at >1% average abundance in 137 geographically diverse cheese microbiomes analyzed in a previous study of rind diversity^18^. These genera are also found in the human gut mycobiome^28^, in the soil^29^, and in marine environments^30^.

**Figure 1:**
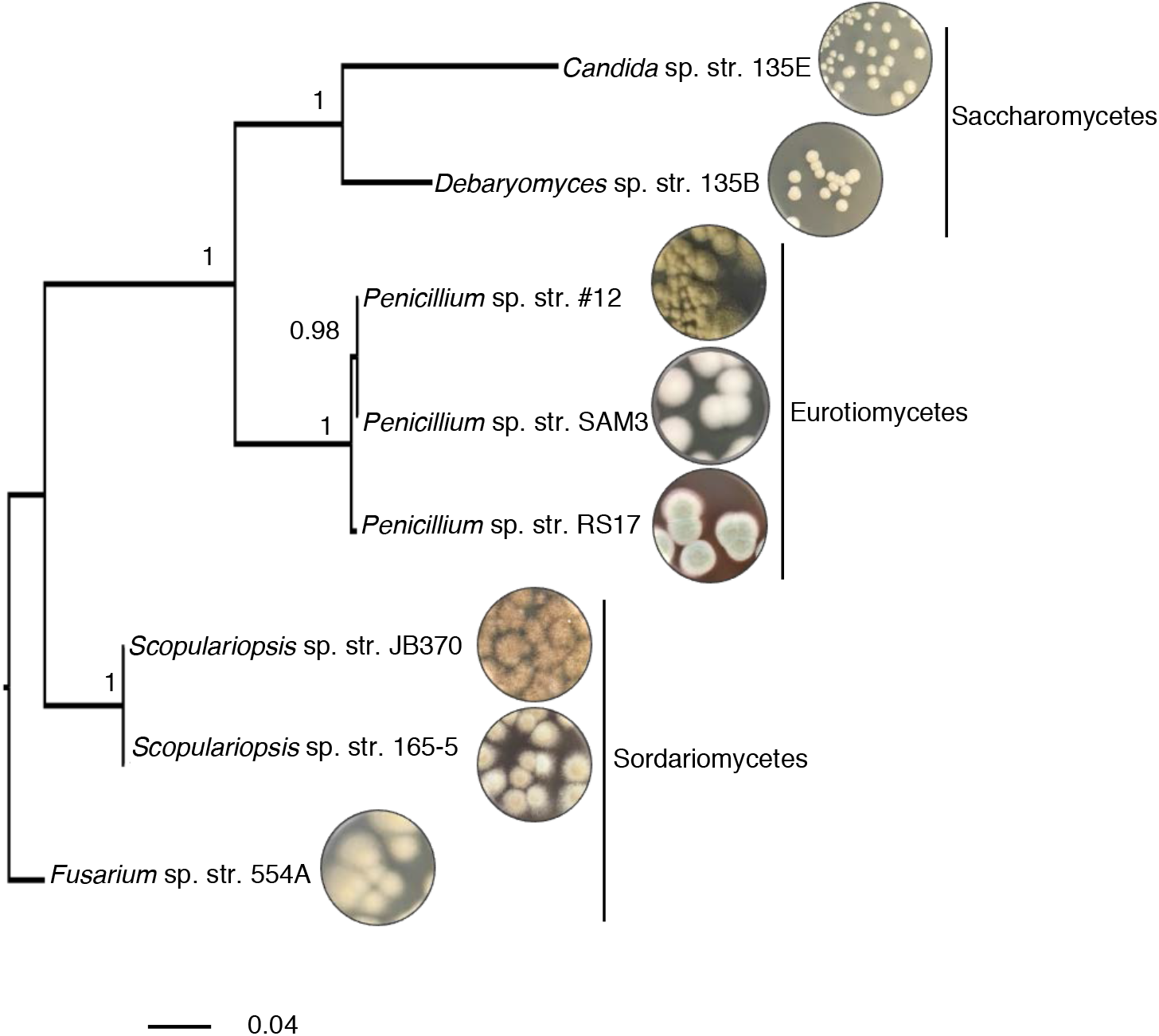
Fungal interaction partners span the phylogenetic and morphological diversity of the cheese ecosystem. Phylogenetic tree based on large subunit rRNA of the cheese fungi used as interaction partners in this study. The tree was built using Bayesian phylogenetic inference with MrBayes^36^ and the Jukes and Cantor substitution model^37^. Branch labels display posterior probability.

The bacterial interaction partners selected were two species of Gammaproteobacteria, *Pseudomonas psychrophila* str. JB418 and *Escherichia coli.* We decided to focus on Proteobacteria for this work, as they are common inhabitants of cheese rind communities from diverse geographic locations and have been shown to be responsive to the presence of fungi in experimental community conditions^18^ (Supplementary Figure 1 and Supplementary Figure 2). *P. psychrophila* is a native and relatively uncharacterized cheese community member isolated from a Robiola due latti cheese rind. *Pseudomonas* species are of interest not only in cheese, but also in human and soil environments^27, 31, 33^. *E. coli* was also included as a bacterial partner in this study to take advantage of the vast genetic resources available for this organism. While *E. coli* is not a common member of cheese rind communities, it is relevant both as a causative agent of foodborne illness in cheese and other foods, and as a commensal member of the human gut microbiome that encounters species from consumed fermented foods^34,35^.

### Characterization of bacterial genes with differential fitness in the presence of fungal partners

Using a pooled library of barcoded transposon-insertion mutants, RB-TnSeq^24^ experiments and analyses generate a fitness value for each gene, reflecting the importance of a gene for survival in the experimental condition. Here, we used RB-TnSeq to identify bacterial mutants that have a differential fitness in the presence of a fungal partner compared to growth alone. To do this, we created a modified experimental and computational pipeline that allowed us to measure and quantitatively compare fitness values across multiple conditions. Previous applications of RB-TnSeq have been designed for intra-condition fitness comparisons, allowing the quantitative comparison of fitness values of different genes within a given condition. However, they were not developed for quantitative inter-condition comparisons, which allow the quantitative comparison of fitness values of the same genes between two different conditions. Specifically, additional normalization steps were needed for inter-condition comparisons. Our updated pipeline includes custom R scripts (available at https://github.com/DuttonLab/RB-TnSeq-Microbial-interactions) which provide step-by-step data visualization to follow the progress of data transformation from raw number of reads to normalized fitness values.

Pooled *P. psychrophila*^21^ or *E. coli*^24^ RB-TnSeq mutant libraries were grown as a lawn for seven days on solid cheese curd agar (CCA) plates^38^ either alone or mixed evenly with one of the eight fungal species (Supplementary Figure 3). Mutant abundances at T0 (inoculation) and day 7 were measured via barcode sequencing and differences in barcode abundance were used to calculate gene fitness values within each condition (Supplementary Data 1 and Supplementary Data 2). To identify genes related to interactions, we looked for mutants whose fitness values were significantly different in a “with fungus” condition versus “alone” condition (p<0.05) (Figure 2). We hereafter refer to these differences as an interaction fitness. In some cases, the presence of a fungus increases the fitness of a mutant (positive interaction fitness), whereas in others the fitness of a mutant is decreased (negative interaction fitness).

**Figure 2:**
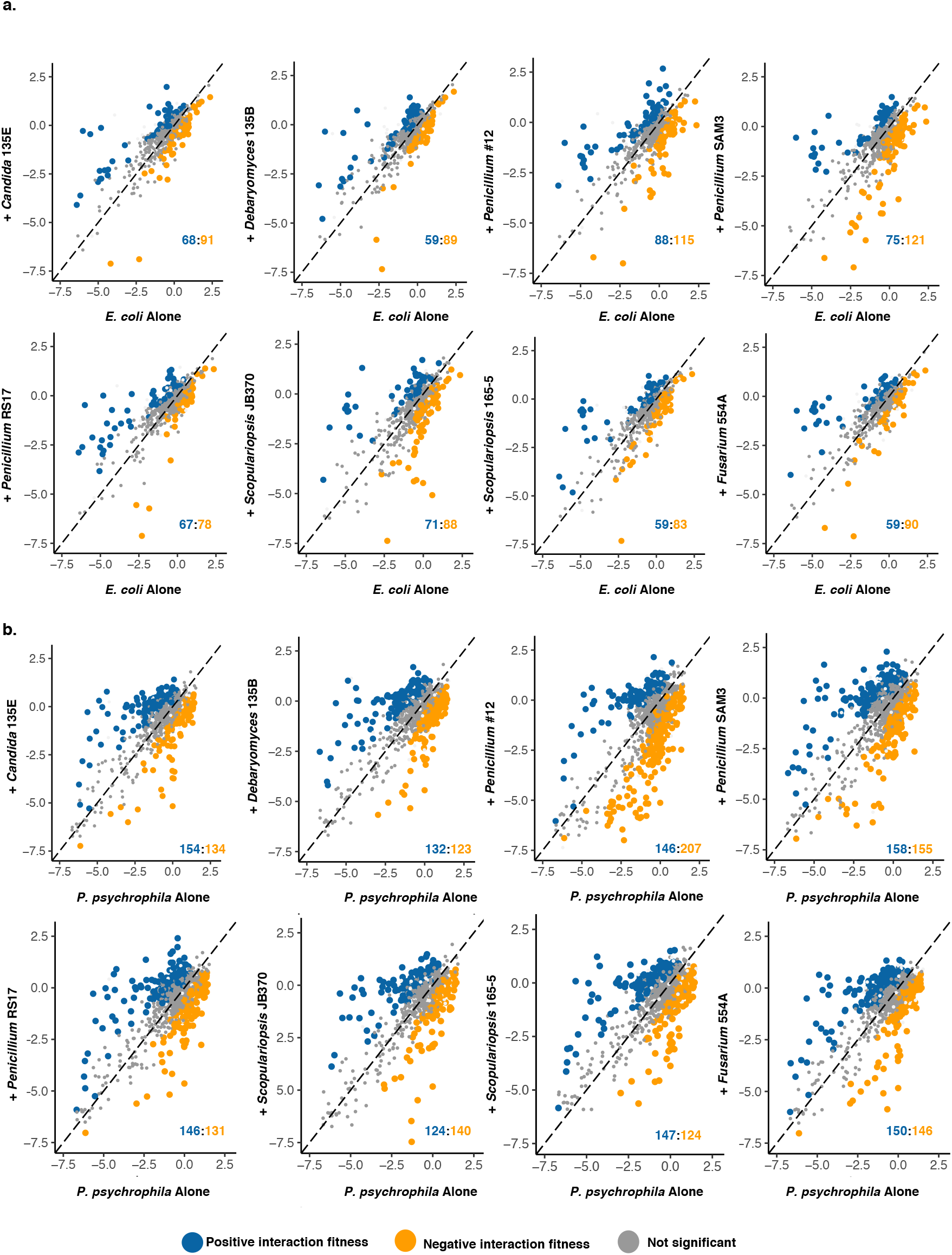
Bacterial genes with significant interaction fitness across fungal partners. Each dot represents a gene, with colored dots indicating genes with a significant interaction fitness. X and Y values indicate gene fitness values in each condition (alone on x-axis versus grown with a fungal partner on y-axis), and the colored numbers in the lower right hand corner indicate how many genes have either positive (blue) or negative (orange) interaction fitness. **a**, *E. coli.* **b**, *P. psychrophila.*

In total, we found 453 *E. coli* and 692 *P. psychrophila* genes whose disruption leads to fitness alteration in the presence of at least one of the fungal partners used in this study (Supplementary Data 3 and Supplementary Data 4). This represents an average of 163 ± 24 *E. coli* genes per fungal condition and 290 ± 32 *P. psychrophila* genes per fungal condition that have interaction fitness (Figure 3a). These gene sets represent around ten percent of the genes of each bacterium. For *E. coli,* interaction fitness values range from −5.66 to 5.72, and for *P. psychrophila,* −6.18 to 6.02. These values are consistent with previously reported fitness values in other experimental systems, where fitness values greater than 2 and less than −2 are generally thought to represent strong fitness effects^24^.

**Figure 3:**
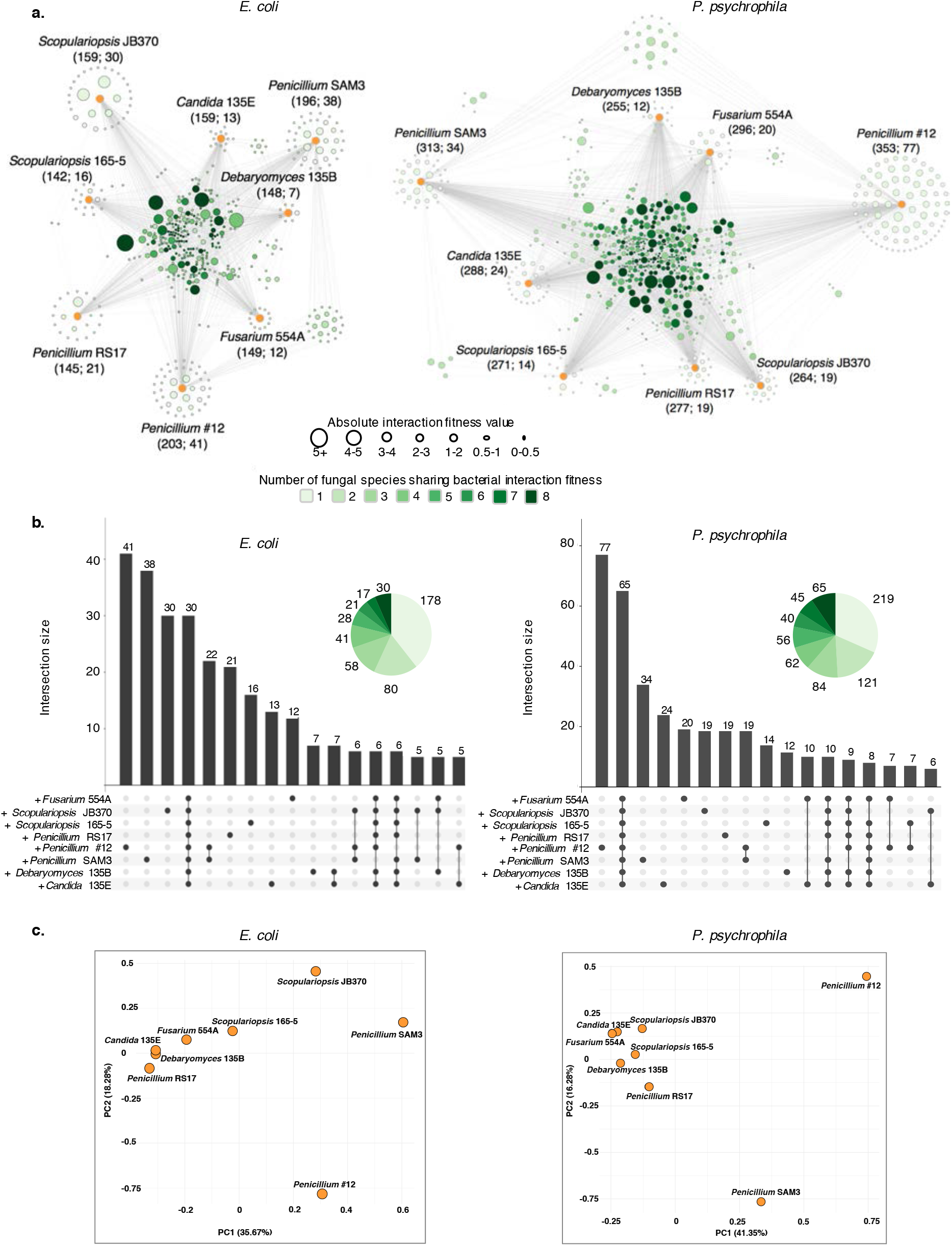
Bacterial genes with differential fitness in the presence of fungi, **a,** Network of *E. coli* (left) or *P. psychrophila* (right) genes with an interaction fitness based on RB-TnSeq. Each orange node represents a fungal partner and is labeled as follows: Fungal partner (Number of genes with interaction fitness; Number of genes with interaction fitness unique to this condition). Each green node represents a bacterial gene. Green nodes are shaded by the number of fungal conditions in which this gene has an interaction fitness as shown in the legend below and are sized by average strength of interaction fitness across partners. **b**, UpSet^39^ plots showing the overlaps, or intersections, of *E. coli* (left) or *P. psychrophila* (right) gene sets with interaction fitness across fungal partners. These UpSet plots are conceptually similar to Venn Diagrams. The connected circles indicate which fungal conditions are included in the intersection, and the size of the intersection (the number of genes that have an interaction fitness in all the highlighted conditions) is displayed in the main bar chart. Intersections <5 genes are not shown. For example, in the *E. coli* panel, 30 genes have an interaction fitness with all partners (all fungi circles are connected), while 22 other genes have an interaction fitness with *Penicillium* sp. str. #12 and with *Penicillium* sp. str. SAM3 (only *Penicillium* sp. str. #12 *and Penicillium* sp. str. SAM3 circles are connected). **c**, PCA of the raw fitness values for all *E. coli* (left) or *P. psychrophila* (right) genes with an interaction fitness in at least one fungal condition.

### Comparison of interaction fitness across fungal partners

To assess the specificity of interactions, we evaluated the intersections of gene sets across the entire set of fungal interaction conditions (Figure 3b, Supplementary Data 5 and Supplementary Data 6). Many genes show differential fitness only in the presence of specific fungal partners. For *E. coli,* 40 percent of the genes with interaction fitness are specific to a single fungus (n=171), and for *P. psychrophila,* 32 percent (n=219). We also identified a number of genes that had interaction fitness across the entire set of partners (n=30 for *E. coli,* and n=65 for *P. psychrophila).* In addition, around 40 percent (n=269) of the interaction-related genes for *P. psychrophila* and 30 percent (n=137) for *E. coli* were common to at least four of the eight fungal interaction conditions. For both *E. coli* and *P. psychrophila,* growth with *Penicillium* sp. str. #12 and *Penicillium* sp. str. SAM3 results in a large number of the same genes with significant interaction fitness (Figure 3a and Figure 3b). These species also do not cluster with the other fungi in Principal Component Analysis (PCA) of the raw fitness values for all *E. coli* (left) or *P. psychrophila* (right) genes having interaction fitness in at least one fungal condition (Fig 3c).

### Mechanisms of fungal impacts on bacterial gene fitness

To identify potential mechanisms underlying bacterial-fungal interactions, we examined the functional distribution of genes associated with interaction fitness. Three functional themes common to multiple fungi used in this study were identified using Clusters of Orthologous Genes (COG) categorization and functional enrichment analysis (Figure 4, Supplementary Data 7 and Supplementary Data 8, Supplementary Figure 4) and analysis of conservation of the effect across fungal species. These interactions include antimicrobial stress on the bacterial cell envelope, competition for biotin, and provision of bioavailable iron.

#### *Penicillium* sp. str. #12 and *Penicillium* sp. str. SAM3 induce bacterial envelope stress

*Penicillium* sp. str. #12 and *Penicillium* sp. str. SAM3 consistently shared impacts on bacterial mutant fitness, as seen by their large number of network connections for both bacteria (Figure 3a and Figure 3b). For *E. coli,* of 116 total genes shared, 22 of these genes are shared only between these two fungi; for *P. psychrophila,* of 187 total genes shared, 19 of these genes are shared only between these two fungi. The gene set shared by these fungi suggests that these two fungal species are producing antibiotic molecules. For *E. coli*, this overlapping gene set includes the genes encoding for the stress protein CspC, beta-lactam antibiotic resistance membrane protein Blr, and the MdtK multidrug efflux pump. Mutants of these genes displayed decreased fitness in the presence of these fungi. This is also supported by the gene set specific to *Penicillium* sp. str. *#12/E.coli,* which includes a regulator of the EmrA multidrug efflux pump and three gene components of the Rcs regulator of capsule synthesis system *(rcsA, rcsC, rcsD).* The Rcs system can respond to damage to the cell envelope and is induced, for example, by membrane-active antimicrobial peptides^40^, peptidoglycan damage caused by inhibition of penicillin-binding proteins^41^, and mutations in lipopolysaccharide biosynthesis genes^42^. Additionally, three genes in the *waa* operon (*waaY, waaP, waaQ)* affecting the modification of the heptose region of the core LPS, which plays a crucial role in membrane stability^43^, have increased fitness in the presence of *Penicillium* sp. str. #12. WaaY activity is dependent on WaaP and WaaQ^43^. Inactivation of *waaY* has previously been shown to increase *E. col’s* resistance to antimicrobial peptide LL-37, due to reduced binding of the peptide to the cell surface^44^, and it has also been shown to increase resistance of *Salmonella typhimurium* to antimicrobial peptides^45^.

When grown in co-culture with *P. psychrophila,* these two fungi induce an interaction fitness for universal stress protein A, whose mRNA is stabilized by the protein encoded by *cspC,* one of the genes with mutants impacted in *E. coli.* With *Penicillium* sp. str. #12, three *P. psychrophila* genes involved in LPS/lipid A/outer membrane biogenesis *(msbA, arnB, waaG)* also had interaction fitness. While the *P. psychrophila* genome is not as well documented as *E. col?s,* these data are consistent with the *E. coli* data and with the production of antibiotics by these fungi.

To investigate fungal antibiotic activity that could be related to the observed changes in bacterial gene fitness, we modified bacterial cytological profiling (BCP) protocols for use on *in vitro* cheese biofilms^46^. This microscopy technique was previously developed to rapidly determine the bacterial cellular pathway targeted by antibiotic compounds. We grew wild type (WT) or Δ*mdtKE. coli* alone or in a mixed biofilm with *Penicillium* sp. str. #12 or *Penicillium* sp. str. SAM3 on CCA plates for seven days. Δ*mdtKE. coli* was chosen due to the RB-TnSeq fitness defect of this mutant specifically in the presence of these *Penicillium.* We performed BCP on bacterial cells either co-cultured in the mixed biofilms or after 2 days of growth on CCA plates with known antibiotic compounds as reference controls (Supplementary Figure 5, Figure 5). Microscopy showed a strong change in cell morphology for both WT and Δ*mdtKE. coli* when grown with *Penicillium* compared to growth alone (Figure 5). When cultured with these fungi, *E coli* cells exhibit a rounded phenotype, consistent with a reduction in cell wall integrity and reminiscent of cells treated with antibiotics that target cell wall biosynthesis such as mecillinam and amoxicillin. Δ*mdtK* cells are strongly affected and have spheroplasted, indicative of the complete loss of structural integrity. Neither of these two fungal strains are known producers of penicillin, and analysis of the *Penicillium* sp. str. #12 draft genome failed to detect penicillin biosynthesis gene clusters^47^. However, these BCP results are consistent with our RB-TnSeq data and suggest that these two fungal strains are inducing stress on the bacterial cell envelope through an undetermined mechanism that may involve novel antimicrobials.

**Figure 4:**
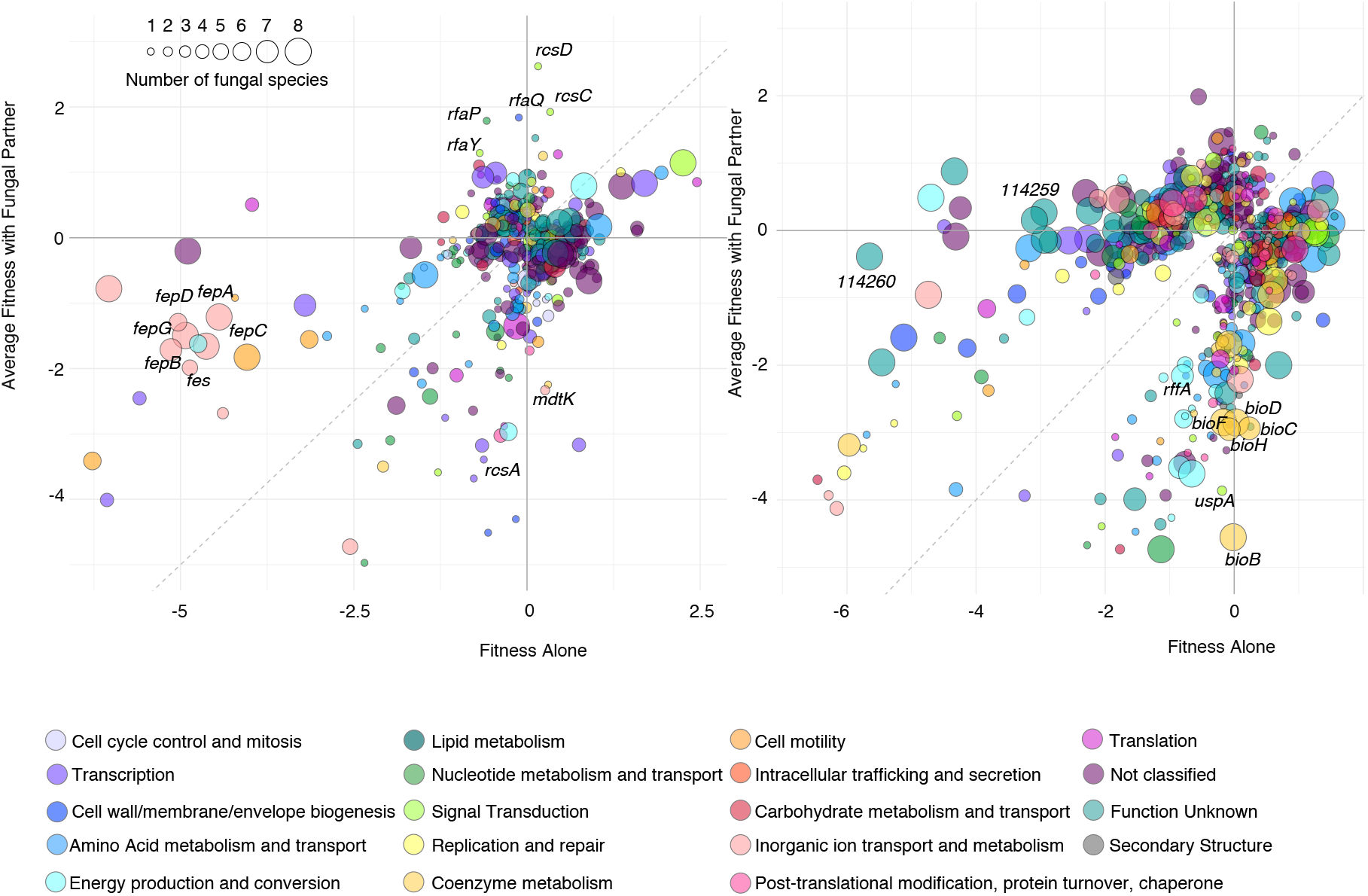
Functional analysis of the bacterial genes associated with interaction fitness. Comparison of *E. coli* (left) or *P. psychrophila* (right) gene fitness values alone compared to fitness values with a fungal partner, colored by COG category and sized by the conservation of effect among fungal partners (1-8 fungal species). Genes discussed in the text are labeled with the gene name.

**Figure 5:**
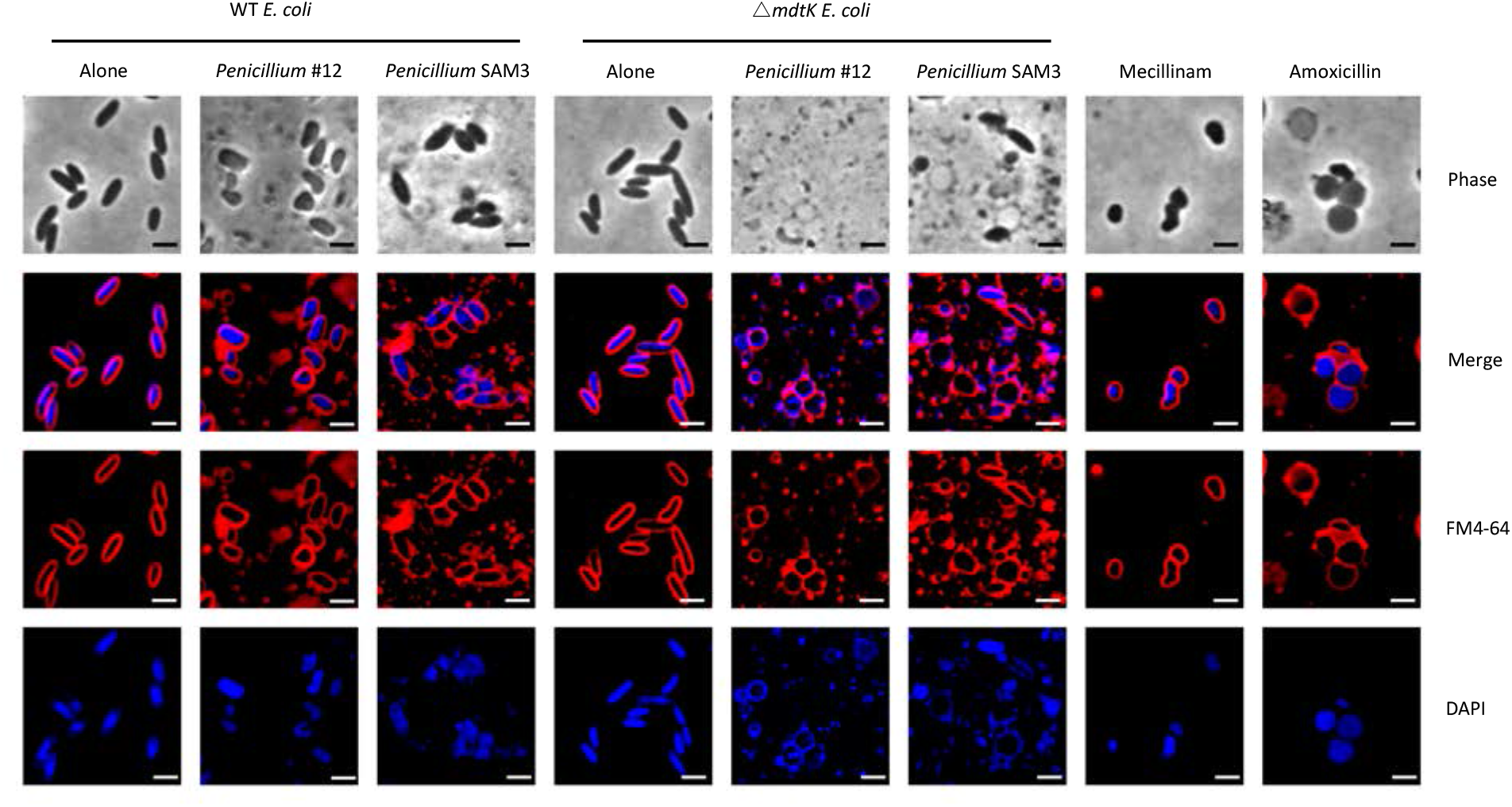
Bacterial cytological profiling of *E. coli* grown in a mixed biofilm with *Penicillium* sp. str. #12 and *Penicillium* sp. str. SAM3 on CCA plates. The phenotype of *E. coli* grown with these fungi is similar to that seen when *E. coli* is exposed to antibiotics targeting cell wall biosynthesis. DAPI dye stains DNA and FM4-64 dye stains bacterial membranes. Scale bars represent 2 μm.

#### Fungi increase bacterial need for biotin biosynthesis

*Pseudomonas* interaction fitness genes suggested competition for environmental biotin between *P. psychrophila* and all of the studied fungal partners. Biotin, found in both milk and cheese^48^, is present in our cheese curd agar medium at 73 nmol/mg and represents an essential cofactor for enzymes involved in key cellular functions like amino acid metabolism and lipid synthesis^49^. Three *P. psychrophila* genes associated with biotin biosynthesis mutants *(bioB, bioD, bioF)* have a negative interaction fitness in the presence of all eight fungi. An additional three genes are associated with a negative interaction fitness in the presence of seven fungi *(bioA, bioC, bioH)* (Figure 4). The biotin biosynthesis pathway was also significantly enriched with fungal partners in functional enrichment analysis of interaction fitness gene sets (Supplementary Data 8). Additionally, differential expression analysis from RNA-Seq of *E. coli* grown either alone or in the presence of *Penicillium* sp. str. #12 showed that *bioA, bioB, bioC, bioD,* and *bioF* were all significantly upregulated in the presence of the fungus with an average fold change of 4.4 (Supplementary Data 9). This highlights an increased need for bacterial biotin synthesis, which again supports that fungi and bacteria are consistently competing for available biotin in the medium, or potentially that bacteria have higher biotin requirements in the presence of fungi. Although some species of fungi are capable of biotin biosynthesis, others are capable of only partial synthesis or are incapable^50^. Notably, this interaction appears to be conserved across both bacteria and all fungi.

#### Fungi increase iron availability for bacterial community members

Because iron is essential for bacterial growth and cheese is an iron-limited environment, with free iron levels measured to be approximately 3 ppm, microbial species growing on cheese require iron chelators with specific transport systems to sustain their growth^21,51–53^. Our RB-TnSeq fitness data revealed that *E. coli* mutants defective in enterobactin transport grow better in the presence of all fungal partners than they do alone (Figure 4, Figure 6a). These genes include members of the *fep* operon *(fepC, fepG, fepA),* which encodes enterobactin transport functions, and *exbD,* which encodes a component of an iron-siderophore transport complex. In most fungal conditions, mutants in an enterobactin esterase encoded by *fes* in addition to iron-siderophore related transport genes *fepB, fepD, tonB* and *exbB* also grow better in the presence of a fungal partner. When grown with any fungal partner, ferric-enterobactin transport is significantly enriched among all the functions associated with genes that have an interaction fitness (Supplementary Data 7).

**Figure 6:**
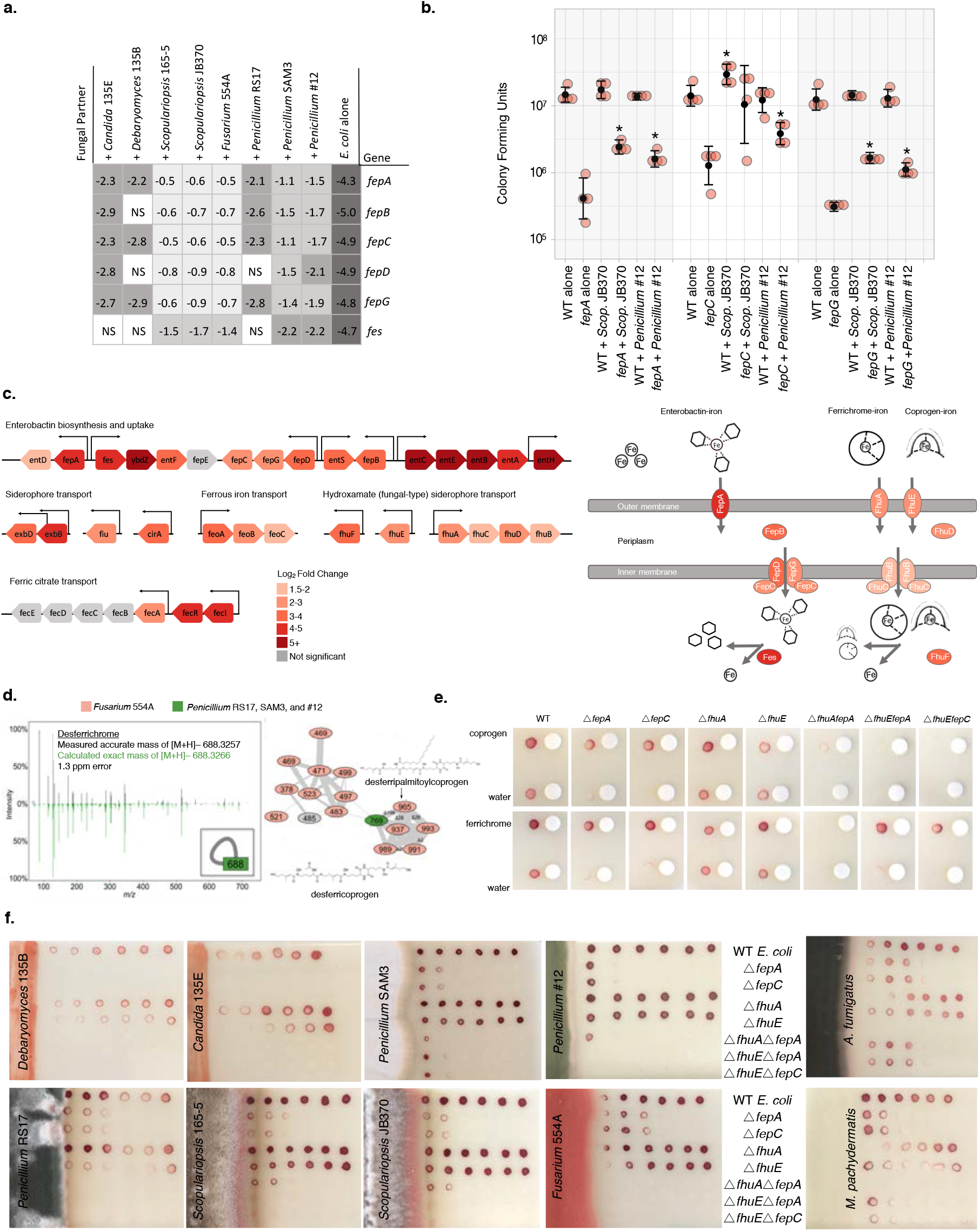
Utilization of fungal siderophores by *E. coli.* **a,** RB-TnSeq fitness values for *fep* operon genes in alone or interactive conditions, showing an increase in fitness in the presence of fungal species. NS-not significant, **b,** Colony forming units of WT and Δ*fep* mutants after 7 days of 1:1 competitive growth on CCA. Competitions were performed either alone or with *Penicillium* sp. str. #12 or *Scopulariopsis* sp. str. JB370. N=4, error bars show standard deviation. Asterisk indicates significantly different growth in the presence of a fungus relative to growth alone (two-sided two-sample equal variance t-test p-value < 0.05). **c**, *E. coli* iron-related genes upregulated in the presence of *Penicillium* sp. str. #12. Significance cutoff made at abs(log_2_(fold-change)) >1.5 and adjusted p-value <0.05. **d**, Fungal siderophores identified by mass spectrometry. Inset on the left shows the node that represents the desferrichrome fragmentation pattern depicted while network on the right represents coprogen-related molecules. Coprogen B and ferrichrome were found by matching fragmentation patterns to library spectra. Both identifications were confirmed using retention time and fragmentation matching to a purchased standard. **e**, Visual assays of Δ*fep* mutant growth with purified siderophores coprogen and ferrichrome. **f,** Visual assays of *E. coli* mutant growth at varying distances from pre-cultured cheese fungi, *A. fumigatus* (soil, human pathogen), and *M. pachydermatis* (skin commensal). Growth performed on CCA containing tetrazolium chloride (red growth indicator).

For *P. psychrophila,* we noticed two genes that have a strong fitness defect alone, but positive interaction fitness when any fungus is present. These genes, Ga0212129_114259 (avg. interaction fitness 3.3) and Ga0212129_114260 (avg. interaction fitness 5.3) are annotated as uncharacterized protein DUF3649 and as an uncharacterized iron-regulated membrane protein. The protein encoded by Ga0212129_114259 is 99% identical to an iron transporter from *Pseudomonas fragi* (NCBI Protein Accession WP_133145017.1). Moreover, immediately upstream of these two genes we find a ferric enterobactin receptor (fepA) and the PfeR-PfeS two-component regulatory system required for production of the ferric enterobactin receptor, suggesting that this region may be involved in siderophore uptake. Again, positive interaction fitness for these likely siderophore-uptake associated genes suggests that possibly more iron is available for *P. psychrophila* when fungi are present.

To validate the RB-TnSeq results related to the Fep system genes, we performed competitive mutant fitness assays with a 1:1 ratio of WT *E. coli* and Δ*fepC*, Δ*fepA*, and Δ*fepG* mutants either alone or in the presence of *Scopulariopsis* sp. str. JB370 or *Penicillium* sp. str. #12. When no fungus is present, there is a clear growth defect for Δ*fep* mutants. After 7 days, both Δ*fepA* and Δ*fepG* mutants grew significantly better with both fungi, whereas there was no difference in wild-type growth with or without fungus. The Δ*fepC* mutant grew significantly better with *Penicillium* sp. str. #12 (Figure 6b).

Additionally, differential expression analysis from RNA-Seq of *E. coli* grown on CCA for three days either alone or in the presence of *Penicillium* sp. str. #12 revealed 34 genes (out of a total of 348 significantly upregulated genes) involved in iron acquisition that are specifically upregulated in the presence of the fungus (Supplementary Data 10). Notably, we find genes associated with the uptake machinery for hydroxamate siderophores (including *fhuA* and *fhuE),* which are commonly produced by fungi. We also observe upregulation of enterobactin biosynthesis and uptake, suggesting that *E. coli,* even in the presence of fungi, still produces and utilizes its native siderophore, enterobactin (Figure 6c).

All filamentous molds in this study, but not yeast, produce siderophores that were detectable by liquid Chrome Azurol S (CAS) assay (Supplementary Figure 6). Given the positive CAS assays, we next sought to identify the siderophores produced by these fungal species. We performed liquid chromatography mass spectrometry (LC-MS and LC-MS/MS) for all fungal species. This metabolomics data showed evidence of the hydroxamate fungal siderophores coprogen and ferrichrome in *Fusarium* and *Penicillium* species (Figure 6d). Although not detected in these extracts, *Scopulariopsis* sp. str. JB370 is predicted to make dimethylcoprogen based on antiSMASH analysis of the draft genome^54^.

We hypothesized that the increased fitness of enterobactin *fep* mutants was due to the uptake of fungal siderophores through an alternate pathway. Indeed, we found that coprogen and ferrichrome can rescue the growth defect of Δ*fepA* and Δ*fepC* on CCA (Figure 6e). The *E. coli* Fhu system has previously been shown to allow uptake of hydroxamate-type siderophores^55–58^.

In *E. coli,* ferrichrome uptake is known to be mediated by the outer membrane receptor FhuA, and coprogen uptake is mediated by the outer membrane receptor FhuE. Mutants of *fhuA* or *fhuE* alone do not show a growth defect on CCA, likely because the enterobactin system is intact. Thus, to specifically examine fungal siderophore uptake, we constructed mutants of *fhuA* or *fhuE* in an enterobactin-uptake defective background. Δ*fepAΔfhuA, ΔfepAΔfhuE*, and Δ*fepCΔfhuE* were grown on CCA in the presence or absence of fungal siderophores (Figure 6e). Combined loss of enterobactin uptake and *fhuA* eliminates the alleviation seen with ferrichrome, whereas loss of either *fhuA* or *fhuE* in the Δ*fepA* background seems to eliminate the alleviation seen with coprogen. This suggests that *E. coli* requires *fhuA* for ferrichrome uptake, and both *fhuA* and *fhuE* for coprogen uptake.

To validate that the fungi used in this study are alleviating *E. coli’s* need for enterobactin through the same mechanism, equal volumes of WT or Δ*fep*/Δ*fhu* mutants were spotted on CCA at varying distances from a pre-cultured fungal front (Figure 6f, Supplementary Figure 7). Growth of Δ*fepA* and Δ*fepC* is restored closest to the fungal fronts of all filamentous molds, but not yeast species. For *Penicillium* sp. str. #12, *Scopulariopsis* sp. str. JB370, and *Scopulariopsis* sp. str. 165-5, this effect is lost in the Δ*fepA*Δ*fhuE* and Δ*fepC* Δ*fhuE* double mutant, suggesting it is *fhuE-dependent.* For *Fusarium* sp. str. 554A and *Penicillium* sp. str. RS17, it is both *fhuE* and *fhuA-dependent.* For *Penicillium* sp. str. SAM3, loss of *fhuA* decreases but does not eliminate alleviation. Thus, we can conclude that near a fungal partner, *E. coli* is likely to use and benefit from fungal hydroxamate siderophores that are taken up by the FhuA and FhuE uptake systems independently of the enterobactin uptake system.

Given that iron limitation is a common challenge across many environments, we wanted to examine whether fungal species from other ecosystems could also be producing siderophores accessible to neighboring bacterial species. We performed similar assays with *Aspergillus fumigatus,* a soil-dwelling filamentous ascomycete that was originally isolated from the lung tissue of a patient who had aspergillosis^59^. This assay was also performed for *Malassezia pachydermatis,* a yeast commensal resident on animal skin. *M. pachydermatis* is also sometimes found on human skin and can act as an opportunistic pathogen; this species has caused bloodstream infections in hospitalized neonates^60 61^. Interestingly, Δ*fepA*Δ*fhuE* and Δ*fepC*Δ*fhuE* mutants are rescued next to *A. fumigatus,* whereas the Δ*fepA*Δ*fhuA* mutant lacking the ferrichrome receptor was not, suggesting that *A. fumigatus* produces a siderophore capable of being imported through FhuA (Figure 6f). *A. fumigatus* is known to produce the extracellular siderophores fusarinine C and triacetylfusarinine C, which are not known to be imported via the Fhu system, and the intracellular siderophore ferricrocin, which is similar to ferrichrome but not expected to be excreted^62^. Ultimately, it is unclear what siderophore is responsible for the effects by *A. fumigatus*. We see a similar effect using *M. pachydermatis,* suggesting that bacteria are able to utilize siderophores from a yeast species using the Fhu system (Figure 6f). *Malassezia restricta* and *Malassezia globosa* have previously been found to possess genes for siderophore biosynthesis^63,64^. We performed AntiSMASH^54^ analysis on a previously published genome of this *Malassezia pachydermatis* strain, and were able to identify a NRPS biosynthetic gene cluster containing a ferrichrome peptide synthetase^54,65,66^. In sum, our results suggest that diverse fungi can reduce bacterial dependence on their own siderophores by secreting xenosiderophores.

### Loss of a fungal secondary metabolite regulator alters the profile of interaction fitness

The cases above show that bacterial gene fitness can be impacted by the production of fungal secondary metabolites, including antimicrobial compounds and siderophores. Previous studies have shown that fungal secondary metabolite production is regulated by the master regulator, LaeA^67^. Loss of LaeA in *Aspergillus* spp., *Fusarium oxysporum,* and *Penicillium chrysogenum* is associated with loss of secondary metabolite production^68,69^. To test the impact of alterations in fungal metabolite production on bacterial gene fitness, we generated a Δ*laeA* mutant in *Penicillium* sp. str. #12.

To assess the impact of the *laeA* knockout on fungal metabolites in *Penicillium* sp. str. #12, we performed RNA-Seq and liquid chromatography with mass spectrometry (LC-MS) on the WT and Δ*laeA* mutant. Fourteen percent of the genome (1925 out of 13261 genes) was differentially expressed between WT and Δ*laeA* (Figure 7a, Supplementary Data 11). This is consistent with previous findings in *A. fumigatus* that LaeA influenced expression of around 10 percent of the fungal genome^67^. GO term enrichment analysis identified melanin, organic hydroxy compound, phenol-containing compound, pigment, sterigmatocystin, depsipeptide, lactone, mycotoxin, and organic heteropentacyclic compound biosynthesis as pathways overrepresented in in the set of 1070 genes more expressed in WT (Supplementary Data 12). Among the genes more expressed in WT, we find four genes in a nonribosomal peptide synthetase cluster region predicted by AntiSMASH^54^; these four genes have homology to *sid2, sidF, sidH,* and *sitT,* genes associated with siderophore biosynthesis and transport in *Aspergillus^62^*. This gene cluster also includes genes with homology to *sidJ* and *mirB,* providing further evidence that this region has a siderophore-related function. Five genes in a predicted gene cluster encoding production of serinocyclin A and four genes in a predicted cluster encoding production of nidulanin A are also more highly expressed in WT than in Δ*laeA*. All together, this confirms that deletion of *laeA* is likely to reduce production of fungal secondary metabolites including siderophores. Among the 855 genes more expressed in Δ*laeA*, GO term enrichment identified amino acid and nucleoside metabolism, fungal cell wall, and ion transport (Supplementary Data 12), highlighting a possible reorganization of *Penicillium* sp. str. #12 metabolism in the absence of *laeA.*

**Figure 7:**
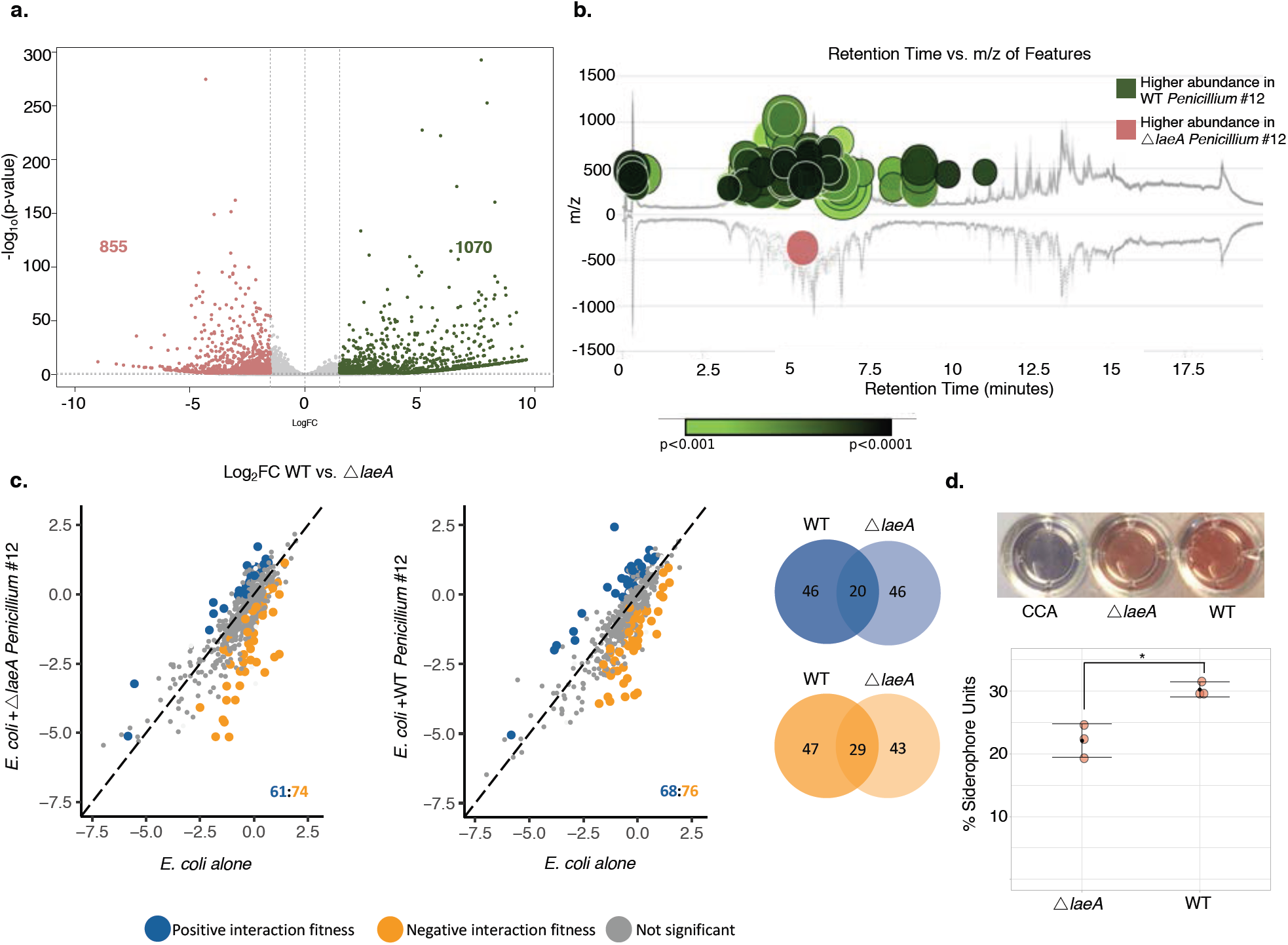
Fungal metabolite production impacts bacterial-fungal interactions, **a,** Differential expression of WT *Penicillium* sp. str. #12 relative to Δ*laeA* after three days of growth on CCA. Labeled on the volcano plot are the number of genes with a log2FC of >1.5 (green) or <-1.5 (red) and adjusted p-value of <0.05. **b,** The metabolomics data analysis platform XCMS^76^ was used to compare features detected by LC-MS analyses of Δ*laeA* and WT *Penicillium* sp. str. #12 extracts. Features of higher abundance in WT relative to Δ*laeA* are depicted as green nodes on the top of the mirror plot and features of lower abundance in WT relative to Δ*laeA* are depicted as red nodes on the bottom. Node radius is proportional to the fold change of the detected features and color intensity is dependent on p-value. The graph displays only those features with a p-value less than or equal to 0.05, fold change higher than or equal to 10, *m/z* between 200 and 2000 Da, and intensity higher than 500. **c,** *E. coli* genes with significant interaction fitness with Δ*laeA* and WT *Penicillium* sp. str. #12. Each dot represents a gene, with colored dots indicating genes with interaction fitness. X and Y values (alone on x-axis and +fungal partner on y-axis) indicate gene fitness values in each condition, and the numbers in the lower right hand corner indicate how many genes have either positive (blue) or negative (orange) interaction fitness. Venn diagrams display the overlap of these gene sets, **d,** Liquid CAS assay of supematants from blank control CCA medium, Δ*laeA,* or WT *Penicillium* sp. str. #12. N=3, error bars show standard deviation from the mean. Asterisk indicates significantly different siderophore production (two-sided two-sample equal variance t-test p-value 0.009).

LC-MS comparison of the two extracts showed differential production of many metabolites, 94 of which have a >10-fold change between the two (Supplementary Data 13). Of these, 93 are less abundant in the Δ*laeA* mutant, which is consistent with the loss of secondary metabolite production in the Δ*laeA* mutant (Figure 7b). Some of these metabolites matched with known secondary metabolites and their analogs based on mass and fragmentation pattern. Namely, atlantinone A and cyclopenol were found to be produced by the WT *Penicillium* sp. str. #12 in >10-fold higher quantity than the Δ*laeA* mutant, and pyripyropene O was putatively identified and produced in >2-fold higher quantity. Cyclopenol is an alkaloid that is described as a mycotoxin in the benzodiazepine class^70^ and serves as an biosynthetic intermediate for viridicatol^71^, which has been described as having antibacterial activity against *Staphylococcus aureus^72^.* Small amounts of viridicatol were indeed found in extracts with high quantities of cyclopenol. Atlantinone A is a meroterpenoid that is derived from the same biosynthetic pathway as other mixed polyketide-terpenoids such as andrastins and citrehybridones produced by various *Penicillium* and *Aspergillus* species^73^. Both pyripyropene O and atlantinone A have tested negative for antimicrobial activity when screened against a panel of bacteria, including an *E. coli* strain^74,75^. The ecological role of these metabolites remains undetermined. In combination with our RNA-Seq analysis, these data highlight an important diminution of secondary metabolite production in the Δ*laeA* strain.

We next performed RB-TnSeq experiments with the *Penicillium* sp. str. #12 Δ*laeA* mutant to determine whether this single fungal mutation leads to changes in the genes needed for bacterial fitness. While some interaction fitnesses are preserved between Δ*laeA* and WT, loss of LaeA appears to significantly change the profile of interactions (Figure 7c). As LaeA is predicted to impact siderophore production, we would expect that *E. coli* enterobactin uptake mutants would not be as strongly impacted by the presence of the Δ*laeA* mutant. Although we see positive interaction fitness of *fepA, fepB, fepC* and *fepG* genes when E. coli is grown with WT *Penicillium* sp. str. #12 relative to *E. coli* growth alone, we do not see positive interaction fitness of *fep* genes with Δ*laeA Penicillium* sp. str. #12 (Supplementary Data 14). Indeed, liquid CAS assays demonstrate that this mutant produces less siderophores than WT on cheese media (Figure 7d). Together, these data support the hypothesis that loss of a fungal secondary metabolite regulator corresponds to changes in bacterial interaction fitness effects.

Our RB-TnSeq results, combined with the antibiotic assay BCP, suggested that antibiotic activity by *Penicillium* sp. str. #12 causes damage to *E. col’s* cell envelope. As the *laeA* mutant should have a decreased ability to produce antimicrobial compounds, we examined whether there were changes in the fitness of genes related to responses to antibiotics. The *mdtK* efflux pump gene, which was previously seen to have negative interaction fitness when grown with WT *Penicillium* sp. str. #12, no longer has an interaction fitness with *ΔlaeA,* suggesting that the Δ*laeA* mutant has decreased antibiotic production. Although the exact nature of the antimicrobials produced by this strain is unknown, these results are consistent with previous studies, which show that LaeA regulates antibiotic production in other fungal species, and with our metabolite analysis, which showed a decrease in production of many metabolites in Δ*laeA*. Overall, these findings suggest that a single fungal gene can strongly impact the bacterial genes needed for fitness and in particular point out that fungal specialized metabolite production may play a large role in determination of bacterial fitness in bacterial-fungal interactions.

## DISCUSSION

Fungi have been shown to strongly impact bacterial neighbors in diverse systems, from soil to polymicrobial infections^77–85^. To characterize the effects of fungi from a simple microbiome on bacteria, we used a high-throughput genetic screen, RB-TnSeq, to identify bacterial genes relevant to fungal interactions. We observed a diversity of interactions, in terms of direction (+/-), strength, and mechanism. These mechanisms include effects on important cellular pathways such as biotin synthesis and antibiotic resistance. Our work demonstrates that there is both a large diversity of bacterial-fungal interactions as well as conservation of key interaction mechanisms across different fungi.

One of the strongest and most widespread bacterial-fungal interactions that we observed suggests that fungal species can dramatically modulate access to iron through the provision of fungal siderophores, such as ferrichrome and coprogen. Although it has long been known that bacteria grown in isolation are able to uptake purified fungal siderophores^55,56^, our results demonstrate that this exchange takes place between bacteria and fungi growing in a biofilm and that this exchange can have impacts on the competitive fitness of bacteria.

Hydroxamate siderophore receptors homologous to FhuE and FhuA are widespread in Proteobacteria, suggesting that this interkingdom siderophore transfer may play an important role in altering metal availability in diverse systems. *Pseudomonas aeruginosa* has previously been shown to produce phenazine metabolites that led to upregulation of extracellular siderophore production by *A. fumigatus^86^.* Fhu uptake systems have also been identified in the Gram-positive bacterial pathogens *Streptococcus agalactiae* and *Listeria monocytogenes,* and growth of non-siderophore producing mutants of *Streptomyces coelicolor* was restored by the presence of siderophores from airborne contaminant fungi^87–90^. Additionally, hydroxamate siderophore uptake systems have been shown to impact *Staphylococcus aureus* fitness in a murine infection model^91^. Another study has shown that the presence of a siderophore-producing, cheese-associated *Scopulariopsis* produced siderophores and caused downregulation of siderophore production by *Staphylococcus equorum^52^. Bacteroides fragilis,* a human gut symbiont, is also able to use ferrichrome to grow in iron-limiting conditions, and *fhu* genes are expressed by *E. coli* in colonic mucus^92,93^. Some fungi consumed as a part of fermented foods have been shown to survive digestive system transit, and fermented foods are known to contain fungal siderophores^94,95^, which could be a source of fungal siderophores in the gut in addition to potential siderophore production by gut-resident species^94–96^. All together, these results suggest that bacterial use of fungal siderophores may be a widespread mechanism of interkingdom interaction. Considering the crucial role of iron acquisition and metabolism in microbial survival, this conserved interaction mechanism could be a determinant in shaping environmental and host-associated microbiomes.

This study provides new insight into the wide range of fungal impacts on bacteria that can occur even in a simple microbiome. Our data also suggest that even among Proteobacteria, fungal impacts may be diverse. Of the 692 *P. psychrophila* genes and 453 *E. coli* genes that had an interaction fitness with at least one fungal partner, only 58 were homologous based on BLAST (Supplementary Data 15). Further, many genes with interaction fitness do not have a known function. We anticipated that by looking for fungal impacts on *E. coli,* we could leverage the vastly superior genetic information available for this species. However, even in this well-characterized organism, 43% of genes identified as having interaction fitness are annotated as hypothetical or putative. For *P. psychrophila,* 29% of genes with interaction fitness are hypothetical proteins. This highlights that many genes involved in interspecies interactions might yet to be characterized, and that studying microbes in the context of their interactions with other species, and not just in monoculture, provides an avenue for uncovering new areas of biology.

## MATERIALS AND METHODS

### Fungal Ribosomal RNA Sequencing

Genomic DNA was extracted with phenol-chloroform (pH 8) from cultures of the eight cheese fungal species used in this study. For each extraction: 125 μL of 425–600 μm acid-washed beads and 125 μL of 150–212 μm acid-washed beads were poured in a screw-capped 2 mL tube. 500 μL of 2X buffer B (200 mM NaCl, 20 mM EDTA) and 210 μL of SDS 20% were added to the tube containing fungal material and 500 μL of Phenol:Chloroform (pH 8). Cells were lysed by vortexing the tubes for 2 min at maximum speed. Aqueous and organic phases were separated by centrifugation at 4°C, 8,000 RPM for 3 min and 450 μL of the aqueous phase (upper phase) were recovered in a 1.5 mL Eppendorf tube. 45 μL of sodium acetate 3M and 450 μL of ice cold isopropanol were added before incubating the tubes at −80°C for 10 min. The tubes were then centrifuged for 5 min at 4°C at 13,000 RPM. The pellet was then washed in 750 μL of 70% ice cold ethanol and re-suspended in 50 μL of DNase/RNase free water. Following DNA extraction, LROR (ACCCGCTGAACTTAAGC) and LR6 (CGCCAGTTCTGCTTACC)^97^ primers were used to amplify the large subunit of the ribosomal RNA and for *Penicillium* species, Bt2a (GGTAACCAAATCGGTGCTGCTTTC) Bt2b (ACCCTCAGTGTAGTGACCCTTGGC)^98^ primers were used to amplify the beta-tubulin gene. PCR was performed in a final volume of 50 μL: 25 μL of Q5 polymerase master mix (New England Biolabs), 2.5 μL of the forward primer at 10 μM, 2.5 μL of the reverse primer at 10 μM, 100 ng of genomic DNA, and water using the following PCR programs: LSU-(i) 98°C – 30 s, (ii) 35 cycles of: 98°C – 10 s; 52°C – 30 s; 72°C – 1.5 min, (iii) 72°C – 5 min; Beta-tubulin-(i) 98°C – 30 s, (ii) 35 cycles of: 98°C – 10 s; 57°C – 30 s; 72°C – 1 min, (iii) 72°C – 5 min. PCR products were purified with the QIAquick PCR purification kit (Qiagen) and sequenced using the forward and reverse primer by Eton Bioscience Inc. (San Diego, USA). Consensus sequences from forward and reverse sequencing reactions were aligned using Geneious version R9 9.1.8 (http://www.geneious.com). The MrBayes^36^ plugin for Geneious was used to build the phylogenetic tree with the following parameters: Substitution model-JC69; Rate variation-gamma; Outgroup-*Fusarium* sp. str. 554A; Gamma categories-4; Chain Length-1100000; Subsampling freq-200; Heated chains-4; Burn-in length-100000; Heated chain temp-0.2; Random seed-1160; Unconstrained branch lengths-1, 0.1, 1, 1. FigTree v1.4.4 was used for visualization (https://github.com/rambaut/figtree/releases).

### Bacterial-Fungal Growth Assays

We aimed to inoculate 60,000 bacterial cells alone or with 6,000 fungal spores per well on 10% CCA medium^38^ adjusted to pH 7 in a 96-well plate. Each bacterial or bacterial-fungal assay was done in triplicate. After 7 days of growth, the entire well was harvested and homogenized in PBS1X-Tween 0.05% prior to dilution and plating on LB with 20 μg/ml cycloheximide (for bacterial counts) or milk plate count agar with 50 μg/ml chloramphenicol (for fungal spore counts). Counts were done at inoculation and after harvest. Final growth counts were then compared in co-culture condition relative to growth alone to identify interaction impacts. Significant growth impacts were determined based on Dunnett’s test^99^, p-value < 0.05. Plots were made with R package ggplot2 3.2.1^100^.

### Microbial Culturing for LC-MS/MS extraction

All fungal cultures were grown on plate count agar milk salt (PCAMS; 1 g/L whole milk powder, 1 g/L dextrose, 2.5 g/L yeast extract, 5 g/L tryptone, 10 g/L sodium chloride, 15 g/L agar). Plates were kept at room temperature and spores were harvested at 7 days of growth (or after sporulation was observed) for subsequent experiments. Spores harvested from fungi were put into PBS and normalized to an OD_600_ of 0.1 for working stock.

### Extraction of cultures

Three biological replicates of each condition were plated (distinct samples) and extracted from solid agar. For extraction from solid agar plates, 5 μL of fungal working stock were spotted onto 10% CCA medium adjusted to pH 7. Following 7 days of growth, agar was removed from the Petri dish and placed into 50 mL falcon tubes. Acetonitrile (10 mL) was added to each tube and all were sonicated for 30 minutes. All falcon tubes were centrifuged and liquid was removed from the solid agar pieces and transferred to 15 mL falcon tubes. The 15 mL falcon tubes containing liquid were then centrifuged and liquid was again removed from any residual solid debris and transferred to glass scintillation vials. These liquid extractions were then dried *in vacuo.* Dried extracts were weighed and diluted with MeOH to obtain 1 mg/mL solutions which were stored at −20°C until analysis via LC-MS/MS.

### LC-MS/MS data collection

High resolution LC-MS and LC-MS/MS data were collected on a Bruker impact II qTOF in positive mode with the detection window set from 50 to 1500 Da, on a 2.1×150mm C18 cortecs UPLC column with a flow rate of 0.5mL/min for a gradient of 10-100% ACN with 0.1% formic acid over 16 minutes. For each sample, 10 μL of a 1mg/mL solution was injected. The ESI conditions were set with the capillary voltage at 4.5 kV. For MS^2^, dynamic exclusion was used, and the top nine precursor ions from each MS^1^ scan were subjected to collision energies scaled according to mass and charge state for a total of nine data dependent MS^2^ events per MS^1^. MS^2^ data for pooled biological replicates is deposited under MassIVE accession number MSV000085070. MS^1^ and MS^2^ data for Δ*laeA* and WT *Penicillium* sp. str. #12 is deposited under MassIVE accession number MSV000085054 and was collected under identical conditions on a Bruker compact qTOF.

### Molecular Networking

For all extractions, all precursor m/z’s that were found in solvent and agar controls (based on both retention time and mass tolerance) were removed prior to input into GNPS using the BLANKA algorithm.^101^ A molecular network (https://gnps.ucsd.edu/ProteoSAFe/status.jsp?task=464b331ef9d54de9957d23b4f9b9db14) was created using the online workflow at GNPS. The data was filtered by removing all MS/MS peaks within +/− 17 Da of the precursor *m/z.* MS/MS spectra were window filtered by choosing only the top 6 peaks in the +/− 50Da window throughout the spectrum. The data was then clustered with MS-Cluster with a parent mass tolerance of .02 Da and a MS/MS fragment ion tolerance of .02 Da to create consensus spectra. Further, consensus spectra that contained less than 2 spectra were discarded. A network was then created where edges were filtered to have a cosine score above 0.7 and more than 6 matched peaks. Further edges between two nodes were kept in the network if and only if each of the nodes appeared in each other’s respective top 10 most similar nodes. The spectra in the network were then searched against GNPS’ spectral libraries. The library spectra were filtered in the same manner as the input data. All matches kept between network spectra and library spectra were required to have a score above 0.7 and at least 6 matched peaks. Solvent and agar control files were also loaded into the networks in order to perform removal based on fragmentation patterns. All nodes with precursor masses less than 200Da were also removed. The extensive background and low *m/z* Da removal was done to more accurately reflect the metabolomic profiles of fungal genera in an attempt to represent only true metabolites. The DEREPLICATOR was used to annotate MS/MS spectra^102,103^. The molecular networks were visualized using Cytoscape software^104^.

### RB-TnSeq Assays

All RB-TnSeq assays were performed on 10% CCA medium adjusted to pH 7. Prior to inoculation, one aliquot of each library was thawed and inoculated into 25 mL of liquid LB-kanamycin (50 μg/mL). Once the culture reached mid-log phase (OD = 0.6–0.8), 5 mL of that pre-culture was pelleted and stored at −80°C for the T0 reference in the fitness analysis. The remaining cells were used to inoculate the fitness assay conditions. For each RB-TnSeq fitness assay, we aimed to inoculate 7,000,000 cells of the bacterial library (on average 50 cells per insertion mutant). For fitness assays including a fungal partner, 700,000 fungal cells were inoculated based on spore counts. Pre-cultured cells were washed in PBS1X-Tween 0.05%, mixed with appropriate volumes of quantified fungal spore stocks, and then inoculated by spreading evenly on a 100 mm petri dish containing 10% CCA medium, pH 7. For each condition, assays were performed in triplicate (3 distinct samples). After seven days, each plate was flooded with 1.5 mL of PBS1X-Tween 0.05% and cells were scraped off, taking care to not disturb the CCA. The liquid was then transferred into a 1.5 mL microfuge tube and cells were pelleted by centrifugation. After removing the supernatant, the cells were washed in 1 mL of RNAprotect solution (Qiagen, Hilden, Germany), pelleted and stored at −80°C until gDNA extraction. Genomic DNA was extracted with phenol-chloroform (pH 8) using the same protocol used for fungal gDNA extraction described above. Samples were stored at −80°C until further analysis.

After gDNA extraction, extracts containing Penicillium sp. str. #12 DNA were purified using the OneStep PCR Inhibitor Removal Kit (Zymo Research, CA, USA). Then, the 98°C BarSeq PCR as described in Wetmore *et al.,* 2015^24^ was used to amplify only the barcoded region of the transposons. PCR was performed in a final volume of 50 μL: 25 μL of Q5 polymerase master mix (New England Biolabs, MA, USA), 10 μL of GC enhancer buffer (New England Biolabs), 2.5 μL of the common reverse primer (BarSeq_P1 – Wetmore et al., 2015) at 10 μM, 2.5 μL of a forward primer from the 96 forward primers (BarSeq_P2_ITXXX) at 10 μM and either 200 ng of gDNA for alone conditions, or 2 μg of gDNA for fungal interaction conditions. For *E. coli* analysis, we performed 84 PCRs (T0 sample and 28 harvest samples in triplicate) involving 28 different multiplexing indexes. For *P. psychrophila* JB418 analysis, we performed 84 PCRs (T0 sample and 28 harvest samples in triplicate) involving 28 different multiplexing indexes. We used the following PCR program: (i) 98°C – 4 min, (ii) 30 cycles of: 98°C – 30 s; 55°C – 30 s; 72°C – 30 s, (iii) 72°C – 5 min. After the PCR, for both *E. coli* and *P. psychrophila,* 10 μL of each of the PCR products were pooled together to create the BarSeq sequencing library and 200 μL of the pooled library were purified using the MinElute purification kit (Qiagen, Germany). The final elution of the BarSeq library was performed in 30 μL in DNase and RNase free water. The BarSeq libraries were then quantified using Qubit dsDNA HS assay kit (Invitrogen, CA, USA) and sequenced on HiSeq4000 (75 bp, single-end reads), by the IGM Genomics Center at the University of California San Diego. The sequencing depth for each condition varied between 6.1 and 11.7 million reads for *E. coli* and 5.8 and 13.3 million reads for *P. psychrophila.*

### RB-TnSeq Data Processing

Custom R scripts were used to determine the average fitness scores for each gene across three RB-TnSeq assay replicates. These scripts are available at https://github.com/DuttonLab/RB-TnSeq-Microbial-interactions. The Readme document provides an in-depth explanation of all data processing steps performed in these scripts. In brief, insertion mutants that did not have a sufficient T0 count (3) in each condition or that are not centrally inserted (10-90% of gene) were removed from analysis. Counts determined by Wetmore *et al*.,^24^ scripts were then normalized to a reference gene *(allB* in *E. coli* and closest protein BLAST match Ga0212129_11710 in *P. psychrophila*) to be able to compare across conditions and to account for differences in sequencing depth. These genes do not have a strong fitness effect in any condition based on former fitness determination developed by Wetmore *et al.,* 2015^24^. Un-normalized strain fitness was then calculated per insertion mutant as the log2 of the ratio of the corrected counts in the condition and the corrected counts in the T0 sample. Un-normalized gene fitness and variance was next calculated by averaging insertion mutants within a gene. These values were then normalized based on the position of the gene along the chromosome, as well as by the mean of the data distribution. These steps were all done on individual replicates. Next, the average gene fitness and associated variance were calculated using the weighted average of the fitness values across the three different replicates. Then, for each condition, fitness values are corrected by the mean of the replicate means (the replicate means being the mean values used to center fitness value within a replicate). Finally, based on the assumption that most genes should have no fitness effect, we corrected gene fitness values in each condition by the mode (peak of the density distribution). Final fitness values were then compared for fungal interaction conditions compared to bacterial alone conditions and comparisons associated with a p-value lower than 5% were considered significant interaction fitness (alpha parameter=0.05 in 2conditions_FitnessComparison.R code). Networks of fitness values were visualized in Cytoscape v. 3.5.1^104^ and PCA plots were made with R package ggplot2 3.2.1^100^ and ggfortify 0.4.7^105^. COG category mapping of *E. coli* and *P. psychrophila* protein sequences was done by eggNOG-mapper v2^106^ and visualized with R package ggplot2 3.2.1^100^.

### Bacterial Cytological Profiling

Approximately 7,000,000 WT *E. coli* K-12 strain BW25113 or Keio collection *mdtK* mutant cells^107^ were inoculated alone or co-inoculated with 700,000 *Penicillium* sp. str. #12 or *Penicillium* sp. str. SAM3 spores on 10% CCA pH 7. After 7 days of growth, 1 mL of T-Base buffer was added to the surface of the biofilms, and biofilms were scraped into the buffer. For co-culture conditions, the sample was filtered through a 0.5 μm filter to specifically remove fungal material. 2 μL of concentrated dye mix (1 μL 1 mg/mL FM4-64, 1 μL 2 mg/mL DAPI in 48 μL T-Base) were added to 20 μL of filtrate. Dye-filtrate mix was spotted onto agarose-LB pads (1% Agarose, 20% LB liquid medium, 80% ddH2O) and imaged by fluorescence and phase contrast. Control compound references on CCA medium were obtained by spotting 30 μL of 5x, 10x, 25x, and 100x MIC dilutions of antibiotics onto quadrants on CCA medium pH 7 plates, allowing to dry, and spread-plating 200 μL of log-phase (OD 0.1) *E. coli* cultures. After two days of growth, cells near the edge of the zone of inhibition on appropriate dilution spots were resuspended in 10 μL of prediluted dye mix (1 μL 1 mg/mL FM4-64, 1 μL 2 mg/mL DAPI in 998 μL T-Base) and spotted onto agarose-LB pads and imaged as described above. Resulting images were deconvolved using Deltavision SoftWorx software (Applied Precision, Inc., WA, USA), analyzed using Fiji^108^, and assembled in Adobe Photoshop (Adobe, CA, USA).

### CCA medium biotin quantification

Biotin quantification of CCA medium was performed on three replicate samples by Creative Proteomics (NY, USA) as follows: 100 mg of each sample was homogenized in water (10 μL/mg) for 1 min three times with the aid of 5-mm metal balls on a MM400 mill mixer. Methanol at 10 μL/mg was then added. Water-soluble vitamins were extracted by vortex-mixing for 2 min and sonication in a water batch for 5 min. After centrifugation, the clear supernatants were cleaned up by solid-phase extraction on a Strata-X (60 mg/mL) cartridge. The eluted fractions containing water-soluble vitamins were collected, pooled and then dried under a gentle nitrogen gas flow in a nitrogen evaporator. The residues were dissolved in 200 μL of 10% methanol. Twenty microliter aliquots were injected to run UPLC-MRM/MS with the use of a C18 UPLC column and with (+) ion detection and (-) ion detection, respectively. Calibration curves were prepared by injection of serially-diluted mixed standard solutions of water-soluble vitamins. Concentrations of detected vitamins were calculated by interpolating the linear calibration curves.

### Δ*fep* mutant competitive growth assays

Approximately 600,000 WT and Δ*fepA*, Δ*fepC,* or Δ*fepG* Keio collection^107^ mutant cells were inoculated at a 1:1 ratio either alone or co-inoculated with approximately 60,000 *Penicillium* sp. str. #12 or *Scopulariopsis* sp. str. JB370 spores on 10% CCA pH 7 in a 96-well plate in four replicates each (4 distinct samples). After seven days of growth, the entire well was harvested and homogenized in PBS1X-Tween 0.05% prior to dilution and plating on LB with 20 μg/ml cycloheximide (total bacterial counts) or with 20 μg/ml cycloheximide and 50 μg/ml kanamycin (bacterial mutant counts). Final growth counts were then compared in co-culture condition relative to growth alone to identify interaction impacts. Significant growth impacts were determined by significantly different growth in the presence of a fungus relative to growth alone based on a two-sided two-sample equal variance t-test p-value < 0.05. Plots made with R package ggplot2 3.2.1^100^.

### RNA-Seq and differential expression analysis of *E. coli* with *Penicillium* sp. str. #12

Approximately 7,000,000 *E. coli* cells were inoculated in triplicate (3 distinct samples) either alone or with approximately 700,000 *Penicillium* sp. str. #12 spores on 10% CCA pH 7. After 3 days, the biofilms were harvested for RNA extraction and washed with 1 mL of RNAprotect. RNA was extracted by a phenol-chloroform extraction (pH 8) using the same extraction protocol as for gDNA extraction. Extractions were then purified with the OneStep PCR Inhibitor Removal Kit (Zymo Research, CA, USA).

Sequencing libraries were prepared as follows. RNA samples were treated with DNase using the ‘Rigorous DNase treatment’ for the Turbo DNA-free kit (AMBION, Life Technologies, Waltham, MA, USA) and RNA concentration was measured by nucleic acid quantification in Epoch Microplate Spectrophotometer (BioTek, Winooski, VT, USA). Transfer RNAs and 5S RNA were then removed using the MEGAclear Kit Purification for Large Scale Transcription Reactions (AMBION, Life Technologies) following manufacturer instructions. Absence of tRNA and 5S RNA was verified by running 100 ng of RNA on a 1.5% agarose gel and RNA concentration was quantified by nucleic acid quantification in Epoch Microplate Spectrophotometer. Also, presence of gDNA was assessed by PCR using universal bacterial 16S PCR primers (Forward primer: AGAGTTTGATCCTGGCTCAG, Reverse Primer: GGTTACCTTGTTACGACTT). The PCR was performed in a final volume of 20 μL: 10 μL of Q5 polymerase master mix (New England Biolabs), 0.5 μL of forward primer 10 uM, 0.5 μL of reverse primer 10 uM and 5 μL of non-diluted RNA. PCR products were run on a 1.7% agarose gel and if gDNA was amplified, another DNase treatment was performed as well as a new verification of absence of gDNA.

Ribosomal RNA depletion was performed using the RiboMinus Transcriptome Isolation Kit (Yeast and Bacteria) for the *E. coli* alone samples and using both the RiboMinus Transcriptome Isolation Kit (Yeast and Bacteria) and the RiboMinus Eukaryote Kit v2 for the mixed *E. coli/Penicillium* sp. str. #12 samples (ThermoFisher Scientific). For the *E. coli* alone samples, each sample was divided into two for treatment and then repooled for RNA recovery with ethanol precipitation. For the *E. coli/Penicillium* sp. str. #12 samples, an equal volume of the eukaryotic probe and RiboMinus Bacterial Probe Mix were used to deplete both bacterial and fungal ribosomal RNA and RNA was recovered with ethanol precipitation. Concentrations after ribosomal RNA depletion were measured using Qubit RNA HS Assay Kits (Invitrogen). The RNA-Seq library was produced using the NEBNext Ultra RNA Library Prep Kit for Illumina for purified mRNA or ribosome-depleted RNA. We prepared a library with a fragment size of 300 nucleotides and used the 10 μM NEBNext Multiplex Oligos for Illumina (Set 1, NEB #E7335) lot 0091412 and the NEBNext multiplex Oligos for Illumina (Set 2, NEB #E7500) lot 0071412. We performed PCR product purification with 0.8X Agencourt AMPure XP Beads. Library samples were quantified with Qubit DNA HS Assay Kits before the quality and fragment size were validated by TapeStation (HiSensD1000 ScreenTape). Library samples were pooled at a concentration of 15 nM each and were sequenced on HiSeq4000 (50 bp, single-end). TapeStation assays and sequencing were performed by the IGM Genomics Center at the University of California San Diego.

Following sequencing, reads were mapped to the concatenated genome of *E. coli* K12 BW25113^109^ and *Penicillium* sp. str. #12 using Geneious version R9 9.1.8 (http://www.geneious.com). Only the reads that uniquely mapped to a single location on the *E. coli* genome section were kept*. E. coli* expression analysis was performed using the following R packages: Rsamtools (R package version 2.0.3), GenomeInfoDb (R package version 1.20.0), GenomicFeatures^110^ (R package version 1.36.4), GenomicAlignments^110^ (R package version 1.20.1), GenomicRanges^110^ (R package version 1.36.1) and DESeq2^111^ (R package version 1.20.1). We followed the workflow described by Love *et al.* and performed the differential expression analysis using the package DESeq2. Differentially expressed genes between two conditions were selected with an adjusted p-value lower than 5% (Benjamini-Hochberg correction for multiple testing) and an absolute log2 of fold change equal to or greater than 1.5.

### Construction of *E. coli* mutants and visual interaction assays

#### Visual assays for hydroxamate siderophore stimulation

Antibiotic assay discs (Whatman) were placed on CCA medium pH 7 with .005% tetrazolium chloride (an indicator of cellular respiration) and 20 μL of water, 10 μM coprogen or ferrichrome (EMC Microcollections GmbH) solutions (in water) were slowly pipetted onto the disc and allowed to absorb. 2.5 μL of 37°C overnight LB cultures of *E. coli* K12 BW25113 WT, Δ*fepA*, Δ*fepC*, Δ*fhuE*, Δ*fhuA*, Δ*fepA*Δ*fhuE*, Δ*fepC*Δ*fhuE*, or Δ*fepA*Δ*fhuA* mutants^107^ were spotted next to the discs. Double mutants were constructed as described below. Plates were left at room temperature until red color developed.

#### Visual assays for fungal stimulation of bacterial mutants

Fungal spores were inoculated on CCA pH 7 with .005% tetrazolium chloride. After fungal growth at room temperature (cheese fungal isolates) or 30°C *(A. fumigatus* and *M. pachydermatis*), 2.5 μL of *E. coli* 37°C overnight LB cultures were spotted at increasing distances from the fungal front. Plates were left at room temperature until red color developed. *A. fumigatus* isolate AF293 was received from Nancy Keller, University of Wisconsin-Madison. *M. pachydermatis* was originally isolated from the ear of a dog in Sweden (ATCC14522).

#### Creation of Δ*fepA*Δ*fhuE* and Δ*fepC* Δ*fhuE*

Chemically competent cells for Δ*fepA* or Δ*fepC* mutants were created. An overnight culture of Δ*fepA* or Δ*fepC* mutants was diluted 1:100 and grown at 37°C until OD 0.4-0.6. The culture was placed on ice for 20 minutes and then centrifuged at 4°C for 10 minutes at 6000 rpm to collect the cells. Supernatant was removed and cells were resuspended in half the previous volume of pre-cooled 0.1M CaCl_2_. After being left on ice for 30 minutes, centrifugation was repeated and supernatant was removed before resuspension in a quarter of the original volume of pre-cooled 0.1M CaCl_2_/15% glycerol. Cells were aliquoted and stored at −80°C until transformation. These cells were transformed with the pKD46 plasmid ^112^, recovered at 30°C and plated on LB plates with 100 μg/mL Ampicillin and grown at 30°C. Overnight cultures were started from individual colonies for creation of electrocompetent cells. Overnight cultures of Δ*fepC*-pkD46 or Δ*fepA-* pkD46 were diluted 1:100 in fresh LB-100 μg/mL Ampicillin and grown at 30°C until an OD of 0.1. 20 μL of fresh 1 M L-arabinose were added and growth was continued at 30°C until OD 0.4-0. 6. Cells were then chilled on ice for 15 minutes and then centrifuged for ten minutes at 4000 rcf 4°C. Cells were resuspended in 1 mL of ice water and centrifuged for ten minutes at 4000 rcf at 4°C. Cells were resuspended in 0.5 mL of ice water and centrifuged for ten minutes at 4000 rcf 4°C. Cells were resuspended in 50 μL of ice water and kept on ice until transformation. The chloramphenicol resistance cassette was amplified from the pKD3 plasmid ^112^ using the custom primers: FhuEcatF (CAGATGGCTGCCTTTTTTACAGGTGTTATTCAGAATTGATACGTGCCGGTAATGGCGCGCCTTACGCCCC) and FhuEcatR (CCTCCTCCGGCATGAGCCTGACGACAACATAAACCAAGAGATTTCAAATGCTGGGCCAACTTTTGGCGAA) and the following PCR conditions (i) 98°C – 30 sec, (ii) 30 cycles of: 98°C – 10 s; 70°C – 20 s; 72°C – 30 s, (iii) 72°C – 5 min. Amplification was performed on 4 ng of pKD3 plasmid using Q5 High-Fidelity 2X Master Mix (New England Biolabs). The PCR product was digested for 1 hour with the restriction enzymes DpnI and ClaI at 37°C and then the PCR product was run on a 1% agarose gel. The PCR product was extracted using the QIAquick Gel Extraction Kit (Qiagen) and then dialyzed for 4 hours with TE buffer. 1.5 μL of dialyzed PCR product was used to transform the electrocompetent Δ*fepC*-pkD46 or Δ*fepA*-pkD46 cells. After 2 hours of recovery in SOC medium with 1mM arabinose at 37°C, the transformation was plated on LB with 50 mg/mL kanamycin and chloramphenicol. Transformants were confirmed to be Δ*fhuE* with Eton Bioscience Inc. sequencing of the chloramphenicol cassette.

#### Creation of Δ*fepA* Δ*fhuA*

Creation was done as with Δ*fepA*Δ*fhuE,* except that the chloramphenicol resistance cassette was amplified from pKD3^112^ using FhuAcatF (ATCATTCTCGTTTACGTTATCATTCACTTT ACATCAGAGATATACCAATGAATGGCGCGCCTTACGCCCCAATGGCGCGCCTTACG CCCC) and FhuAcatR (GCACGGAAATCCGTGCCCCAAAAGAGAAATTAGAAACGGAAGGTTGCGGTCTGGGCCAACTTTTGGCGAACTGGGCCAACTTTTGGCGAA) custom primers.

### *Penicillium* sp. str. #12 genome sequencing, assembly, and annotation

Genomic DNA was extracted from *Penicillium* sp. str. #12 using the genomic DNA extraction protocol described above. High molecular weight DNA (average 16 Kb) was sequenced on the Oxford Nanopore MinION with a R.9.5 flow cell using 1D^2^ sequencing adaptors from kit SQK-LSK308 (Oxford Nanopore Technologies, Oxford, United Kingdom). Raw data was basecalled using guppy 3.3.0 (Oxford Nanopore Technologies, Oxford, United Kingdom)(guppy_basecaller-config dna_r9.5_450bps.cfg --fast5_out) for 1D basecalls and these were used to also obtain higher accuracy 1D^2 basecalls (guppy_basecaller_1d2 -i 1Dbasecall/workspace/ --config dba_r9.5_450bps_1d2_raw.cfg --index_file 1Dbasecall/sequencing_summary.txt) These reads were assembled by canu 1.8^113^ and polished by racon 1.4.3^114^ four times and by pilon 1.23^115^ once. The final assembly is 38 Mbp and consists of 52 contigs.

*Penicillium* sp. str. #12 genome annotations were obtained by combining genomic and transcriptomic information from RNA-Seq. To obtain the gene expression profile of *Penicillium* sp. str. #12, approximately 700,000 WT *Penicillium* sp. str. #12 spores were inoculated in triplicate on 10% CCA pH 7. After 3 days, the biofilms were harvested for RNA extraction and washed with 1mL of RNAprotect. RNA was extracted and RNA-Seq libraries were prepared as described above with the following modification: Ribosomal RNA depletion was performed using the RiboMinus Eukaryote Kit v1 and RNA was recovered with ethanol precipitation. After sequencing, the RNA-Seq reads from these *Penicillium* sp. str. #12 alone cultures were concatenated with RNA-Seq reads from the previously described *E. coli/Penicillium* sp. str. #12 co-culture conditions that uniquely mapped to a single location on the *Penicillium* sp. str. #12 genome. The full set of transcriptomic reads were then used as input into the FunGAP annotation pipeline and 77 million of these reads mapped^116^. This pipeline predicted 13261 protein-coding genes in the *Penicillium* sp. str. #12 genome. Interproscan^117^ was used within the FunGAP pipeline for function prediction of genes. This Whole Genome Shotgun project has been deposited at DDBJ/ENA/GenBank under the accession JAASRZ000000000. The version described in this paper is version JAASRZ010000000.

### Creation and confirmation of *laeA* deletion in *Penicillium* sp. str. #12

Deletion cassette design strategy: In order to knockout *laeA* in *Penicillium* sp. str. #12, the isolate was first screened for hygromycin and phleomycin resistance. *Penicillium* sp. str. #12 showed a confirmed sensitivity to both antibiotics. A three round PCR deletion strategy was used to replace the *laeA* ORF with the *hph* gene, whose expression confers selection on hygromycin^118^. The schematic representation of the *laeA* gene replacement with the hph gene is depicted in Supplementary Figure 8. The deletion cassette (5’flank-hph-3’flank) was constructed using three sequential PCR reactions. In the first PCR round, about 1 Kb genomic sequence flanking either the 5’ or 3’ end of the *laeA* ORF were amplified using the primer sets P12_KOlaeA_5’ F (CTCCGTTGGGCCCTCAC) and 5’R (GCAATTTAACTGTGATAAACTACCGCATTAAAGCTGTTGATATCGGCAATCAATCA ATG) or P12_KOlaeA_3’F (GGTGGGCCTTGACATGTGCAGCCGGTGGAGCGGCGCCTGGTGAATCCTACCCACATGG) and 3’R (CGTTGGGAGGAAAAGCTTCTGCG) respectively. The *hph* gene was amplified from plasmid pUCH2-8 using primers hph_F (AGCTTTAATGCGGTAGTTTATCACAG) and hph_R (CTCCACCGGCTGCACATGTC). A second PCR reaction was performed to assemble by homologous recombination the three individual fragments from the first round PCR. The deletion cassettes were finally amplified using the nested primer set, P12_KOlaeA_NestedF (CAGACGGTCCGCATCCCG) and P12_KOlaeA_NestedR (GGTCCAGGTGCAGTAGTACTG).

Fungal transformation: To generate the deletion strains, a protoplast-mediated transformation protocol was employed. Briefly, 109 fresh spores were cultured in 500 mL of liquid minimal medium (LMM) for 12 h under 25°C and 280 rpm. Newly born hyphae were harvested by centrifugation at 8000 rpm for 15 min and hydrolyzed in a mixture of 30 mg Lysing Enzyme from Trichoderma (Sigma-Aldrich) and 20 mg Yatalase (Fisher Scientific) in 10 mL of Osmotic Medium (1.2 M MgSO_4_, 10 mM NaPB, pH 5.8). The quality of protoplast was monitored under the microscope after four hours of shaking under 28 °C and 80 rpm. The protoplast mixture was later overlaid with 10 mL of trapping buffer (0.6 M sorbitol, 100 mM Tris-HCl pH 7.0) and centrifuged for 15 min under 4°C and 5000 rpm. Protoplasts were collected from the interface, overlaid with an equal volume of STC (1.2 M sorbitol, 10 mM Tris-HCL pH 7.5,10 mM CaCl_2_) and decanted by centrifugation at 6000 rpm for 8 min. The protoplast pellet was resuspended in 500 μL STC and used for transformation. After 5 days of incubation at 25 °C, colonies grown on stabilized minimal medium (SMM) plates supplemented with hygromycin were subjected to a second round of selection on hygromycin plates. In total, 25 hygromycin-resistant transformants were isolated after a rapid screening procedure on SMM supplemented with hygromycin. Single-spored transformants were later tested for proper homologous recombination at the ORF locus by PCR and Southern blot analysis.

Gene-deletion strain confirmation: The correct replacement of the *laeA* with the *hph* gene was first verified by PCR analysis of genomic DNA derived from the transformant strains using primer set P12_laeA_F (CACAATGGCTGAACACTCTCGG) and P12_laeA_R (GGGATATGGAGCATCGAAGTTGC) that amplify the *laeA* ORF. About 12% (3/25) of the monoconidial lines generated from primary transformants of *Penicillium* sp. str. #12 were PCR-positive for the absence of the *laeA* ORF. The positive deletion strains were further checked for a single insertion of the deletion cassette by Southern blot analysis and revealed single-site integration of the deletion cassette in one transformant (Supplementary Figure 8). Probes corresponding to the 5’ and 3’ flanks of the *laeA* gene in each strain were labeled using [α32P] dCTP (PerkinElmer, USA) following the manufacturer’s instructions.

### RNA-Seq analysis of WT and Δ*laeA Penicillium* sp. str. #12

To characterize the effect of the *laeA* deletion on the *Penicillium* sp. str. #12 gene expression profile, we performed RNA-Seq analysis for Δ*laeA Penicillium* sp. str. #12. As for WT *Penicillium* sp. str. #12, 700,000 Δ*laeA Penicillium* sp. str. #12 spores were inoculated in triplicate (3 distinct samples) on 10% CCA pH 7 and biofilms were harvested after 3 days. Harvest, RNA extraction and library preparation were performed identically to WT *Penicillium* sp. str. #12. Then, *Penicillium* sp. str. #12 and Δ*laeA* differential expression analysis was performed as described for *E. coli/Penicillium* sp. str. #12 above. To look for enrichment of functions in the set of differentially expressed genes, we input the protein sequences of the genes into the gene-list enrichment function of KOBAS 3.0^119^. Sequences were searched against the Gene Ontology (GO) database^120,121^ using *A. fumigatus* as a reference for GO assignment before conducting a hypergeometric test with Benjamini-Hochberg correction. Functions with a corrected p-value <.05 were considered enriched.

### Data Availability

Sequence data that support the findings of this study (RB-TnSeq, RNA-Seq) have been deposited in the NCBI SRA database with BioProject PRJNA624168. Mass spectrometry data is available in the MassIVE database under accession numbers MSV000085070 and MSV000085054. The GNPS molecular network is available at https://gnps.ucsd.edu/ProteoSAFe/status.jsp?task=464b331ef9d54de9957d23b4f9b9db14. The Whole Genome Shotgun project for *Penicillium* sp. str. #12 has been deposited at DDBJ/ENA/GenBank under the accession JAASRZ000000000 in BioProject PRJNA612335.

The version described in this paper is version JAASRZ010000000. In addition to these sources, source data used to create figures 2,3,4, 6, and 7 is available in the Supplementary Data provided with the paper.

### Code Availability

The R scripts developed for processing RB-TnSeq data described in this manuscript are available at https://github.com/DuttonLab/RB-TnSeq-Microbial-interactions along with usage instructions. The perl scripts needed for initial processing of RB-TnSeq data published in Wetmore *et al.* 2015^24^ are available at https://bitbucket.org/berkeleylab/feba/src/master/.

### Availability of biological materials

All unique materials, including described fungal strains isolated from cheese, the *P. psychrophila* JB418 strain and RB-TnSeq library, *laeA Penicillium* sp. str. #12 deletion mutant, and *E. coli* siderophore uptake double mutants, are readily available from the authors upon request. The *E. coli* RB-TnSeq library and Keio strains can be requested from the groups that created these resources (PMID references provided). *Penicillium* sp. str. SAM3 is commercially available from Danisco.

| Resource Sources |  |  |  |
| --- | --- | --- | --- |
| Resource | Source or reference | Identifiers | Additional information |
| Keio collection, strain background *Escherichia coli* K12 | PMID: 16738554 | CGSC,<br>RRID:SCR_002303 | Collection of 3,818 *E. coli* knockout strains |
| Keio_ML9 library, strain background *Escherichia coli* K12 | PMID: 25968644 |  | RB-TnSeq library of *E. coli* K12 BW25113 (152,018 pooled insertion mutants) |
| JB418_ECP1 library, strain background *Pseudomonas psychrophila* JB418 | PMID: 30211673 |  | RB-TnSeq library generated in the *P. psychrophila* JB418 strain isolated from cheese (272,329 pooled insertion mutants) |
| Keio ME9062, strain background *Escherichia coli* K12 | PMID: 16738554 | CGSC#: 7636 | Parent strain of the Keio collection mutants |
| *Penicillium* sp. str. SAM3 | Danisco - CHOOZIT | PC SAM 3 LYO 10D | Industrial starter for cheese production |
| *Pseudomonas psychrophila* JB418 | PMID: 30211673 |  | Strain isolated from Robiola due latti cheese |
| *Candida* sp. str. 135E | This paper |  | Strain isolated from cheese |
| *Debaryomyces* sp. str. 135B | This paper |  | Strain isolated from cheese |
| *Penicillium* sp. str. #12 | This paper |  | Strain isolated from cheese |
| *Penicillium* sp. str. RS17 | This paper |  | Strain isolated from cheese |
| *Scopulariopsis* sp. str. JB370 | This paper |  | Strain isolated from cheese |
| *Scopulariopsis* sp. str. 165-5 | This paper |  | Strain isolated from cheese |
| *Fusarium* sp. str. 554A | This paper |  | Strain isolated from cheese |
| *Aspergillus fumigatus* str. AF293 | Nancy Keller,<br>University of Wisconsin-Madison |  |  |
| *Malassezia pachydermatis* str. CBS 1879 | ATCC14522 |  |  |

## ACKNOWLEDGMENTS

The authors would like to thank: the Arkin lab and the Deutschbauer lab at UC-Berkeley for the *E. coli* Keio_ML9 library, Kristen Jepsen at the IGM Genomics Center at the University of California San Diego for assistance with sequencing, Dr. Sergey Kryazhimskiy (UCSD) for his input on RB-TnSeq data processing, Cong Dinh (UCSD) for assistance with fungal genome assembly, William Bushnell for assistance with fitness validation experiments, Dr. Sinem Beyhan for advice on fungal genome annotation, Lisa Marotz (UCSD) for assistance with non-cheese yeasts, and members of the Dutton lab, especially Brooke Anderson, for constructive comments on the manuscript.

## SUPPLEMENTAL FIGURE AND TABLE LEGENDS

**Supplementary Figure 1: Impacts of fungal species on bacterial growth after 7 days of coculture on cheese curd agar, pH 7.** CFU: colony forming units. N=3, error bars show standard deviation and black point is the mean. * represents a significant difference in bacterial growth in the presence of the fungal partner relative to alone (two-sided Dunnett’s test, p-value <.05).

**Supplementary Figure 2: Impacts of bacterial species on fungal growth after 7 days of coculture on cheese curd agar, pH 7.** For filamentous fungi, spore counts were used as a proxy for fungal CFUs. N=3, error bars show standard deviation and black point is the mean. * represents a significant difference in fungal growth in the presence of the bacterial partner relative to alone (two-sided Dunnett’s test, p-value <.05).

**Supplementary Figure 3: RB-TnSeq assay.** Characterized pooled bacterial mutant libraries were grown in a biofilm either alone or in a mixed biofilm with a fungal partner. After seven days of growth, mutant abundances were compared to the starting library abundances for each condition. Changes in barcode abundances were used to calculate gene fitness values. Genes with fitness values that differed significantly between co-culture and alone conditions (significant interaction fitness) were identified as potentially relevant to fungal interaction.

**Supplementary Figure 4: COG categories of genes with interaction fitness.** Number of genes with interaction fitness falling into each COG category for *E. coli* (left) or *P. psychrophila* (right).

**Supplementary Figure 5: Bacterial Cytological Profiling of Δ*tolCE.coli* treated with known antibiotic compounds on cheese curd agar.** DAPI dye stains DNA and FM4-64 dye stains bacterial membranes. SYTOX green stains nucleic acids but cannot penetrate live cells. Scale bars represent 2 μm.

**Supplementary Figure 6: Siderophore production by filamentous molds.** Liquid CAS assay was performed on filtered and concentrated fungal supernatants from three replicates grown in liquid cheese for 12 days. Row A) 1-3: Liquid cheese control 4-6: *Penicillium* SAM3. Row B) 1-3: *Debaryomyces* 135B 4-6: *Penicillium* #12. Row C) 1-3: *Candida* 135E. 4-6: *Penicillium* RS17. Row D) 1-3: *Scopulariopsis* 165-5 Row E) 1-3: *Scopulariopsis* JB370. Row F) 1-3: *Fusarium* 554A. % Siderophore units calculated as [(A_r_ – A_s_)/(A_r_)]*100, where A_r_ is the absorbance of the cheese curd agar supernatant blank and A_s_ is the absorbance of the sample. N=3, error bars show standard deviation and black point is the mean.

**Supplementary Figure 7: Fitness defect of *fep* mutants on iron-limiting CCA.** Visual assays of *E. coli* mutant growth spotted alone on CCA pH 7.

**Supplementary Figure 8. Deletion of *laeA* gene in *Penicillium* sp. str. #12.** A. Schematic representation of the genetic construct for *laeA* deletion in *Penicillium* sp. str. #12. The construct is constituted of the *hph* gene conferring resistance to hygromycin. The positions of the restriction enzyme cutting sites are shown on the map. B. Southern blot analyses of genomic DNA from the WT and the Δ*laeA* strains. Ten micrograms of total DNA from each strain was digested with the appropriate enzymes and subjected to Southern blot analysis using respectively the 5’ flank fragment (blue) and the 3’fragment (grey) as probes. The 1 Kb DNA ladder from New England Biolabs was used to determine the size of the expected bands. The blot image was cropped to place the confirmed mutant adjacent to the positive control. The transformants that were confirmed to not have the correct insertion were not included in the figure.

**Supplementary Data 1: RB-TnSeq fitness values for *E. coli* grown with fungal partners compared to alone.**

**Supplementary Data 2: RB-TnSeq fitness values for *P. psychrophila* grown with fungal partners compared to alone.**

**Supplementary Data 3: Genes with significant interaction fitness for *E. coli* grown with fungal partners.**

**Supplementary Data 4: Genes with significant interaction fitness for *P. psychrophila* grown with fungal partners.**

**Supplementary Data 5: Intersection lists for *E. coli* genes with interaction fitness across all conditions.**

**Supplementary Data 6: Intersection lists for *P. psychrophila* genes with interaction fitness across all conditions.**

**Supplementary Data 7: Functional enrichment results for *E. coli* genes with interaction fitness across all conditions.**

**Supplementary Data 8: Functional enrichment results for *P. psychrophila* genes with interaction fitness across all conditions.**

**Supplementary Data 9: RNA-Seq differential expression analysis for *E. coli* grown with *Penicillium* sp. str. #12 *(E. coli* perspective, alone condition as reference).**

**Supplementary Data 10: Iron-related genes that are differentially expressed by *E. coli* when grown with *Penicillium* sp. str. #12 relative to *E. coli* growth alone.**

**Supplementary Data 11: Metabolite production by Δ*laeA* and WT *Penicillium* sp. str. #12. Δ*laeA* used as the reference condition.**

**Supplementary Data 12: RB-TnSeq fitness values for *E. coli* grown with Δ*laeA* or WT *Penicillium* sp. str. #12 compared to alone.**

**Supplementary Data 13: RNA-Seq differential expression analysis for Δ*laeA* vs. WT *Penicillium* sp. str. #12 (WT used as the reference condition).**

**Supplementary Data 14: Functional enrichment results for genes differentially expressed in Δ*laeA* vs. WT *Penicillium* sp. str. #12.**

**Supplementary Data 15: Overlap of *E. coli* or *P. psychrophila* genes with interaction fitness.**

