## Supplementary Figure 1 for "From iron to antibiotics: Identification of conserved bacterial-fungal interactions across diverse partners"

### Slide 1
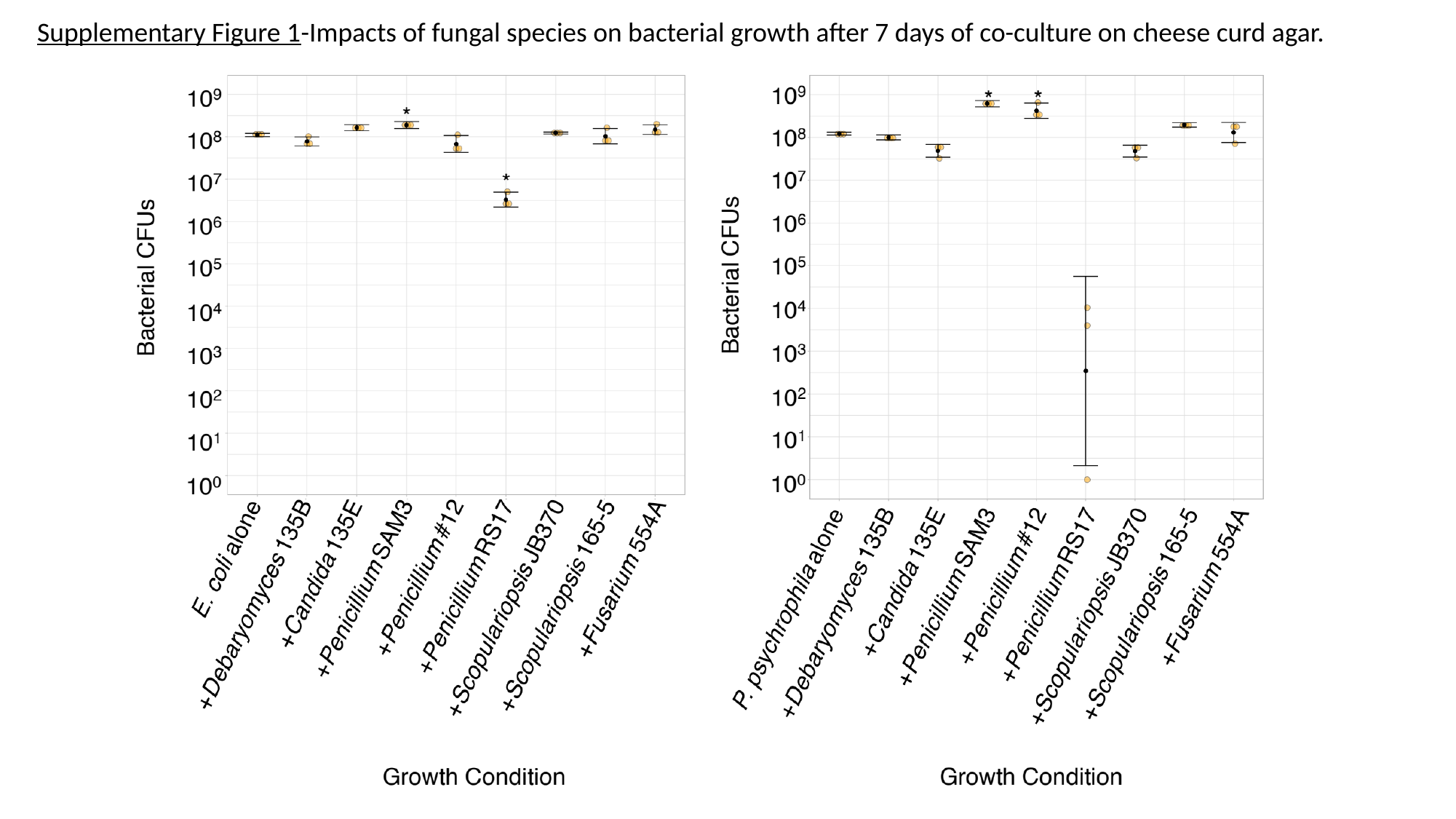

Supplementary Figure 1-Impacts of fungal species on bacterial growth after 7 days of co-culture on cheese curd agar.
