## Supplementary figures and images for "From iron to antibiotics: Identification of conserved bacterial-fungal interactions across diverse partners"

### Supplementary Figure 3

#### Slide 1
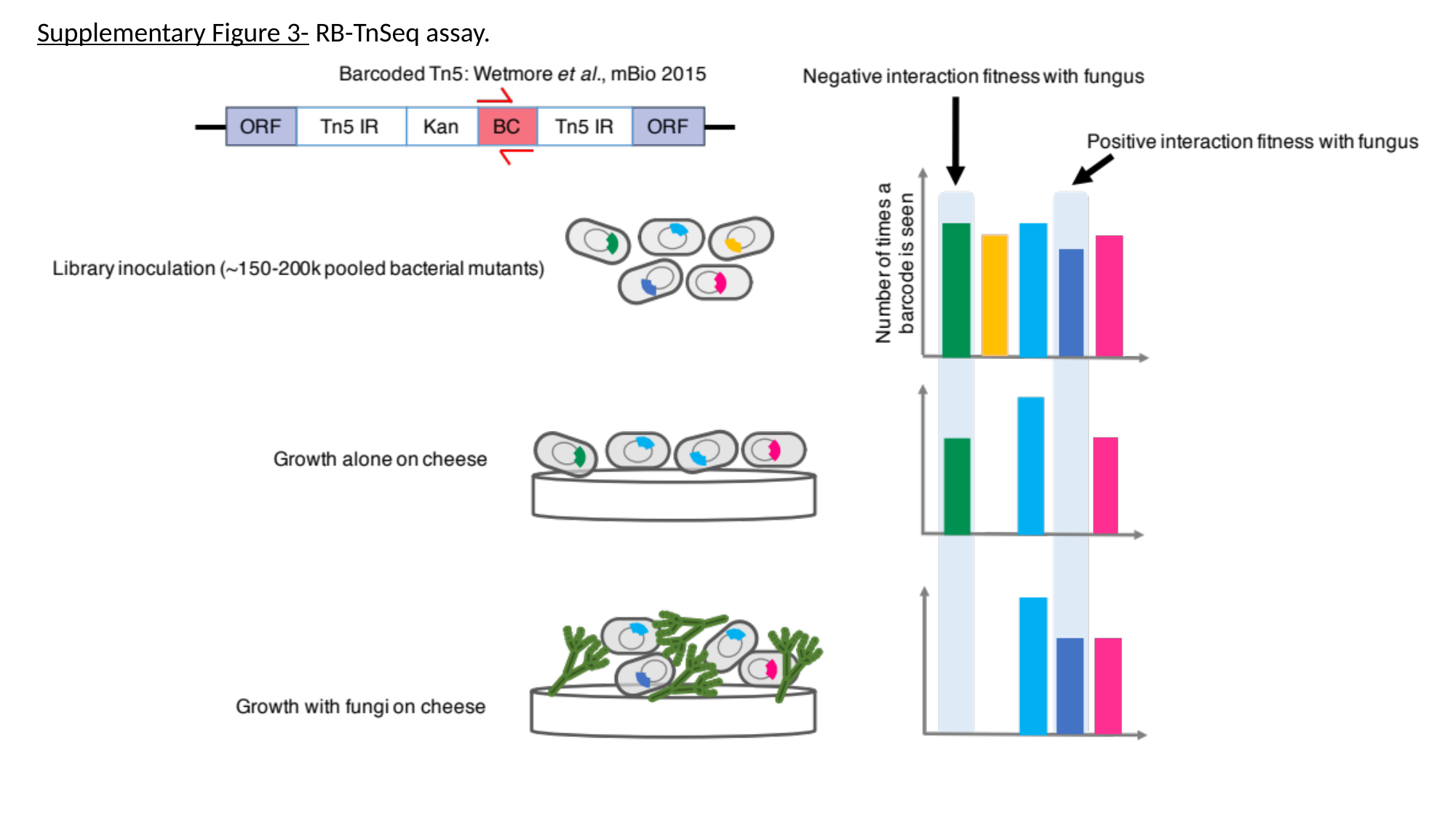

Supplementary Figure 3- RB-TnSeq assay.

### Supplementary Figure 4

#### Slide 1
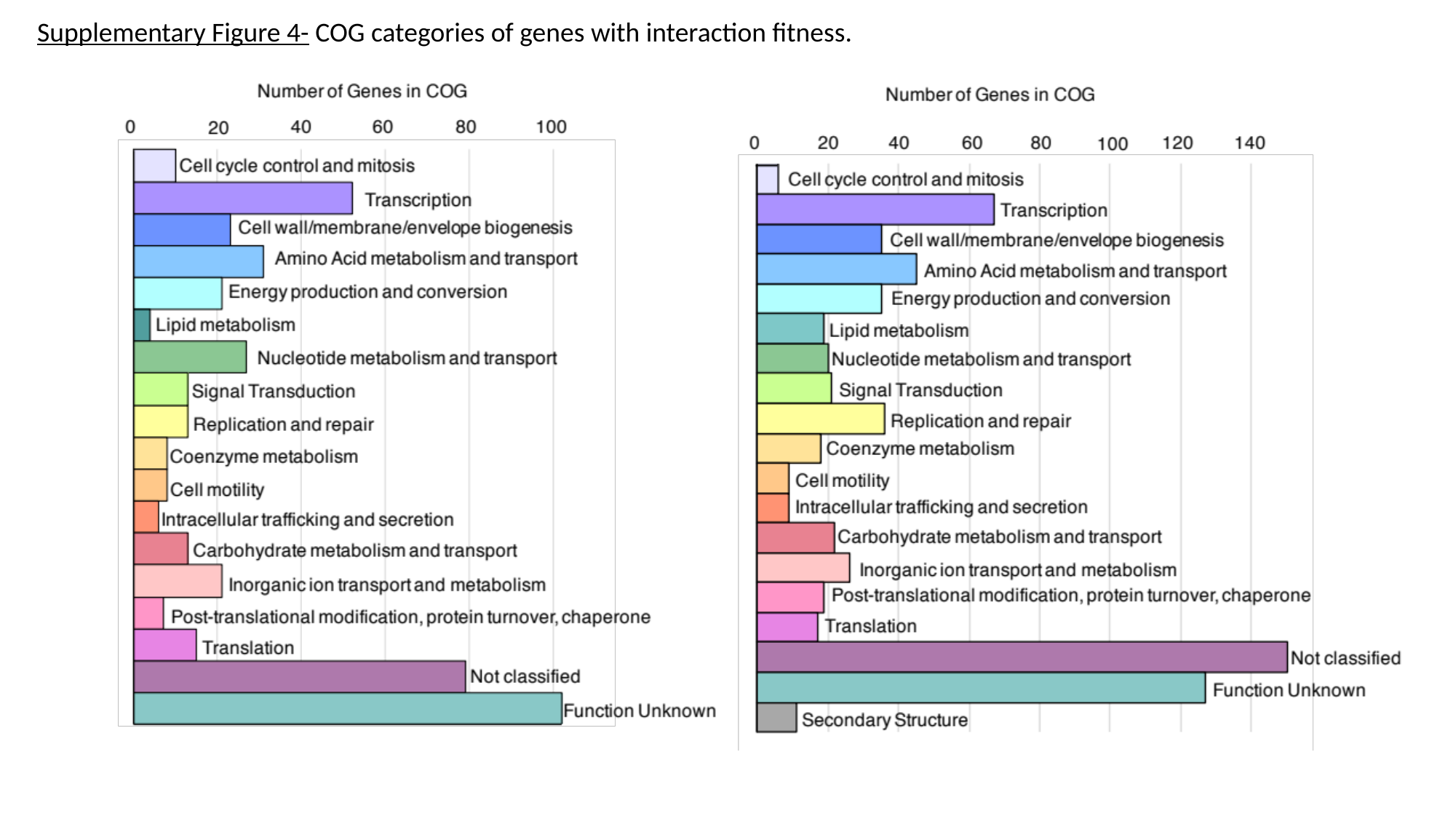

Supplementary Figure 4- COG categories of genes with interaction fitness.

### Supplementary Figure 6

#### Slide 1
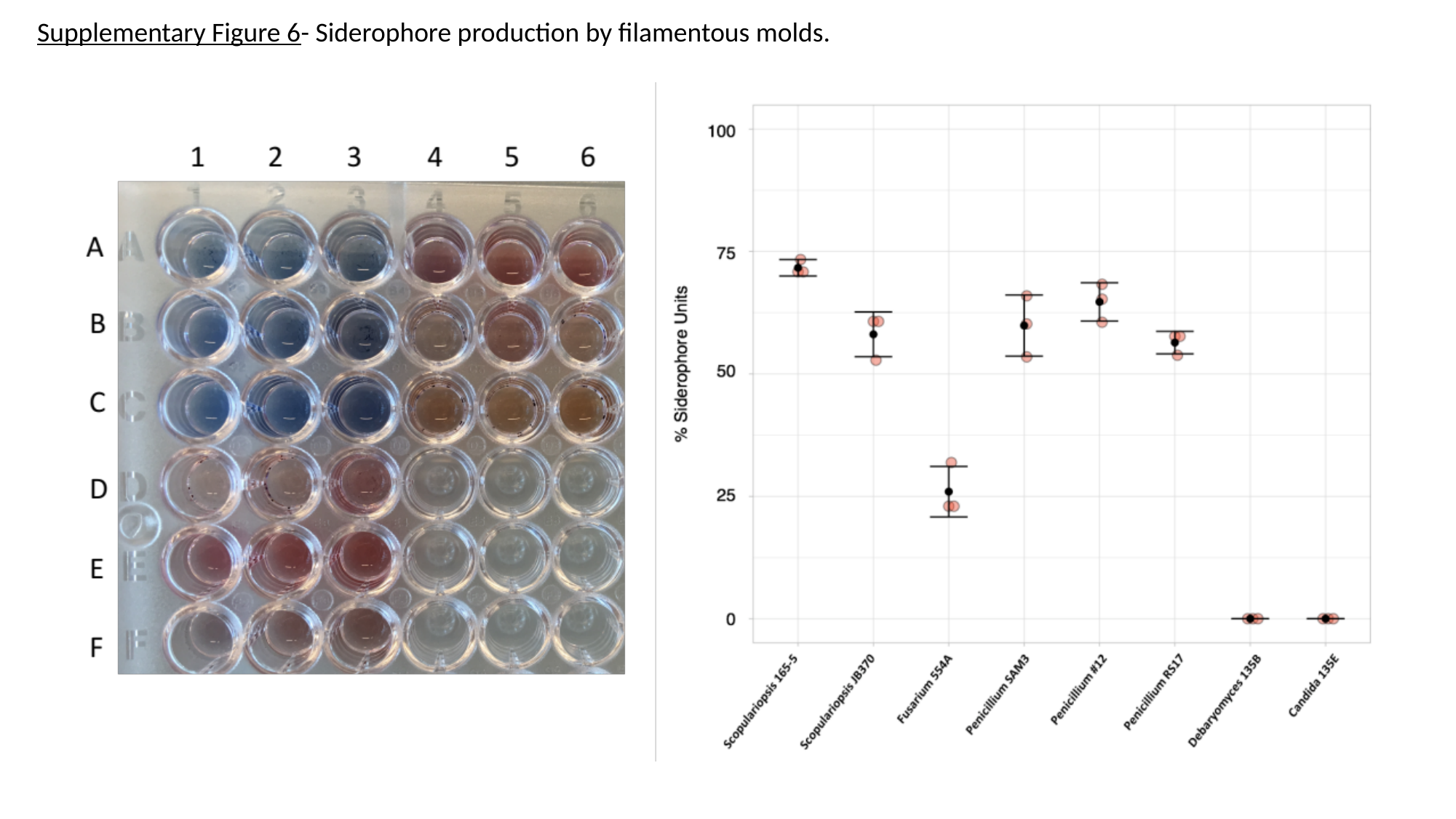

Supplementary Figure 6- Siderophore production by filamentous molds.

### Supplementary Figure 7

#### Slide 1
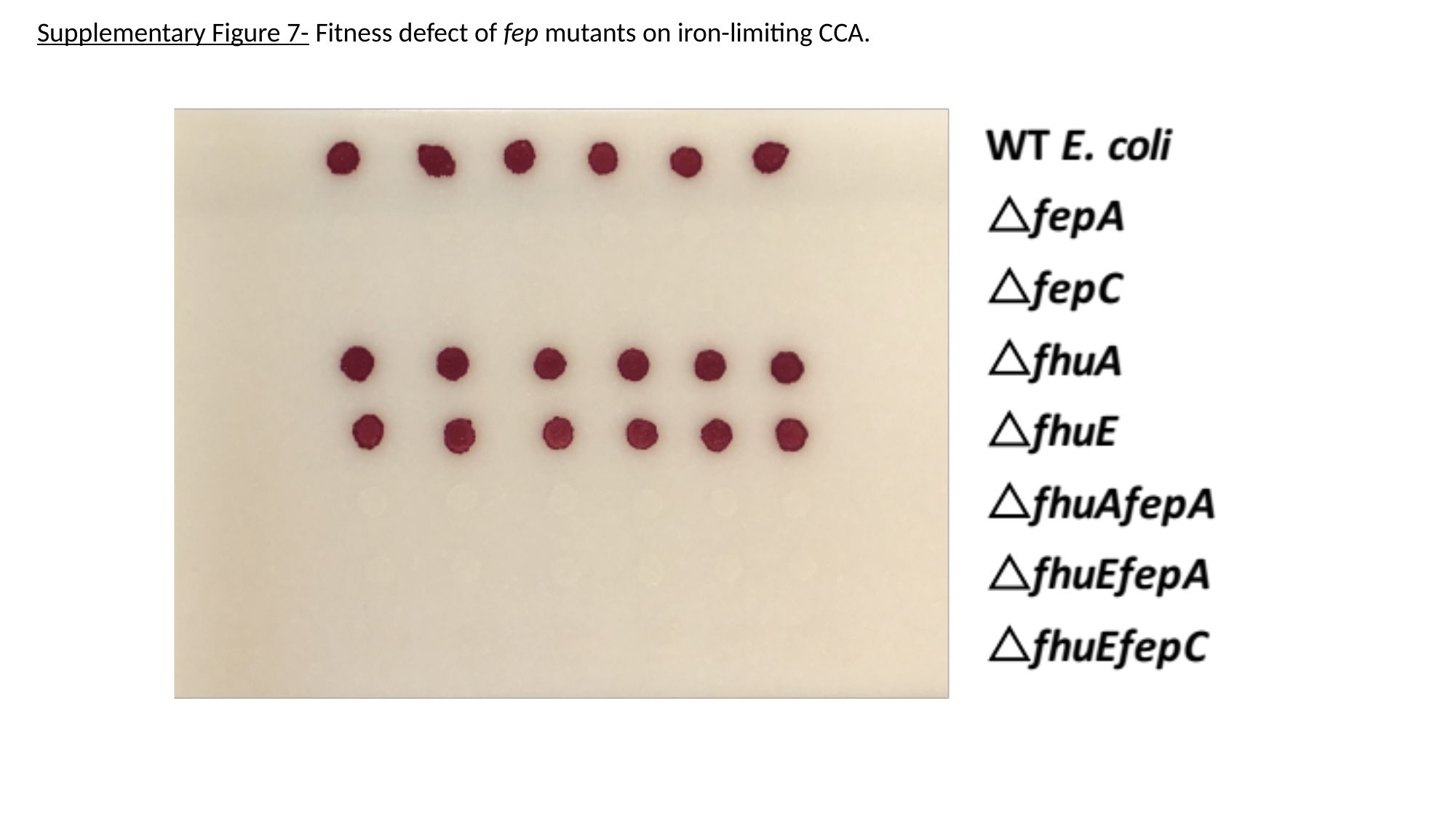

Supplementary Figure 7- Fitness defect of fep mutants on iron-limiting CCA.

### Supplementary Figure 8

#### Slide 1
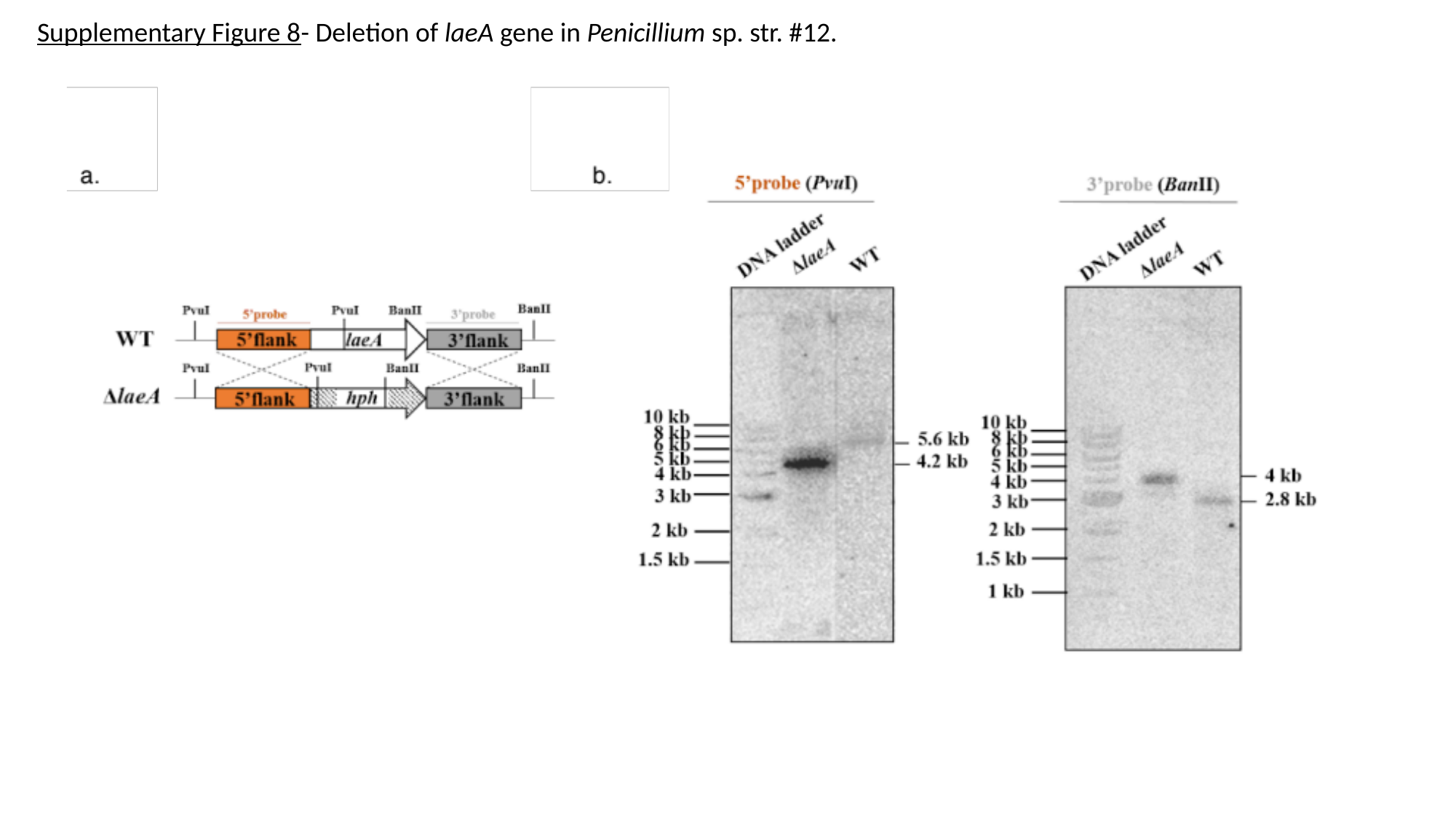

Supplementary Figure 8- Deletion of laeA gene in Penicillium sp. str. #12.
