## Supplementary Figure 5 for "From iron to antibiotics: Identification of conserved bacterial-fungal interactions across diverse partners"

### Slide 1
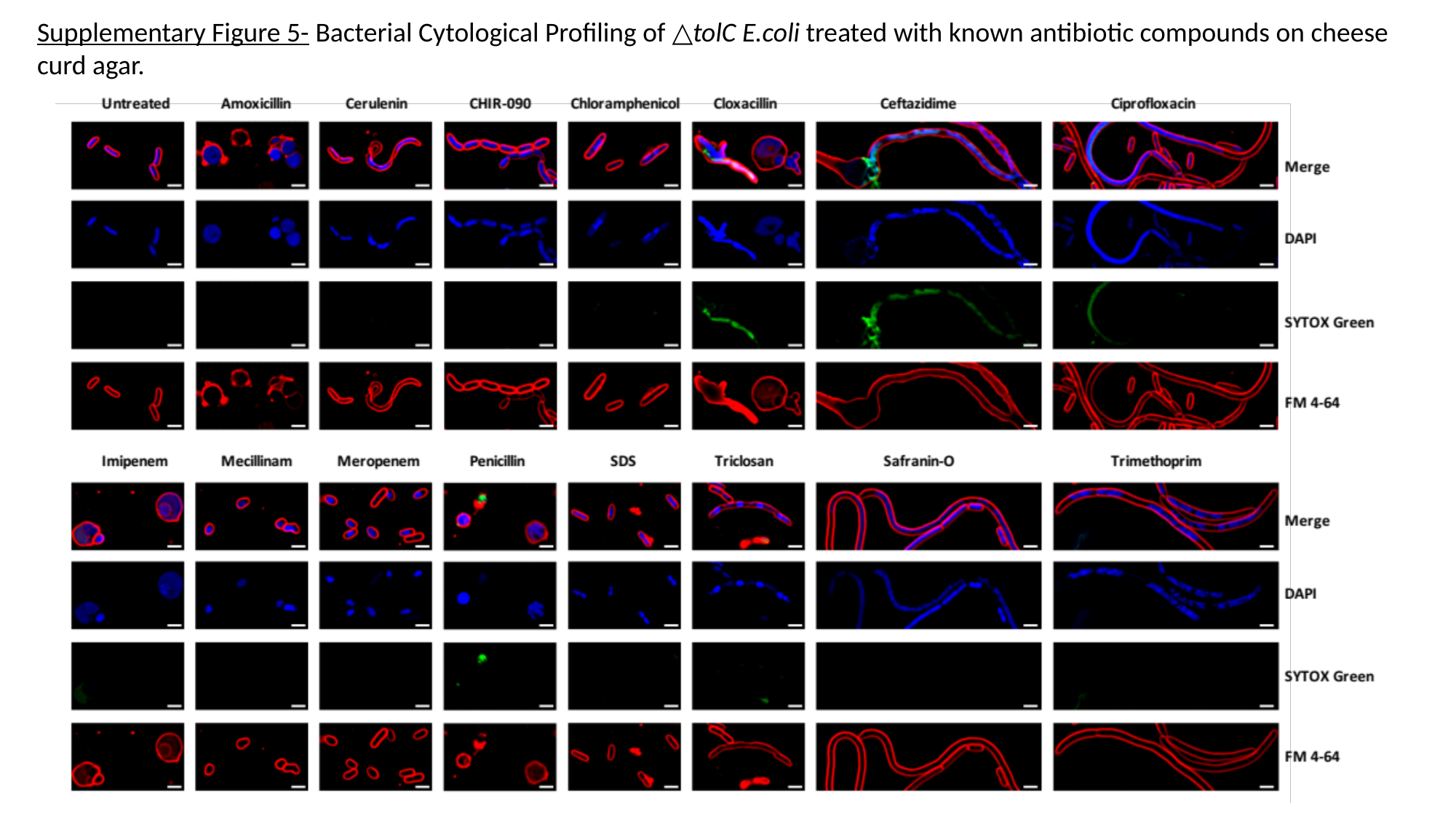

Supplementary Figure 5- Bacterial Cytological Profiling of △tolC E.coli treated with known antibiotic compounds on cheese curd agar.
